# A combined flow injection/reversed phase chromatography – high resolution mass spectrometry workflow for accurate absolute lipid quantification with ^13^C- internal standards

**DOI:** 10.1101/2020.11.04.367987

**Authors:** Harald Schoeny, Evelyn Rampler, Yasin El Abiead, Felina Hildebrand, Olivia Zach, Gerrit Hermann, Gunda Koellensperger

## Abstract

We propose a fully automated novel workflow for lipidomics based on flow injection-followed by liquid chromatography high resolution mass spectrometry (FI/LC-HRMS). The workflow combined in-depth characterization of the lipidome achieved via reversed phase LC-HRMS with absolute quantification as obtained by a high number of lipid species-specific- and/or retention time (RT) matched/class-specific calibrants. The lipidome of ^13^C labelled yeast (LILY) provided a cost efficient, large panel of internal standards covering triacylglycerols (TG), steryl esters (SE), free fatty acids (FA), diacylglycerols (DG), sterols (ST), ceramides (Cer), hexosyl ceramides (HexCer), phosphatidylglycerols (PG), phosphatidylethanolamines (PE), phosphatidic acids (PA), cardiolipins (CL), phosphatidylinositols (PI), phosphatidylserines (PS), phosphatidylcholines (PC), lysophosphatidylcholines (LPC) and lysophosphatidylethanolamines (LPE). In order to exploit the full potential of isotopically enriched biomass, LILY was absolutely quantified on demand via reversed isotope dilution analysis using FI-HRMS. Subsequent LC-HRMS analysis integrated different calibration strategies including lipid species-specific standards for >90 lipids. Extensive measures on quality control allowed to rank the calibration strategies and to automatically selected the calibration strategy of highest metrological order for the respective lipid species. Overall, the workflow enabled a streamlined analysis pipeline (identification and quantification in separate analytical runs) and provided validation tools together with absolute concentration values for >350 lipids in human plasma on a species level with an analytical run-time of less than 25 min per sample.

**TOC:** 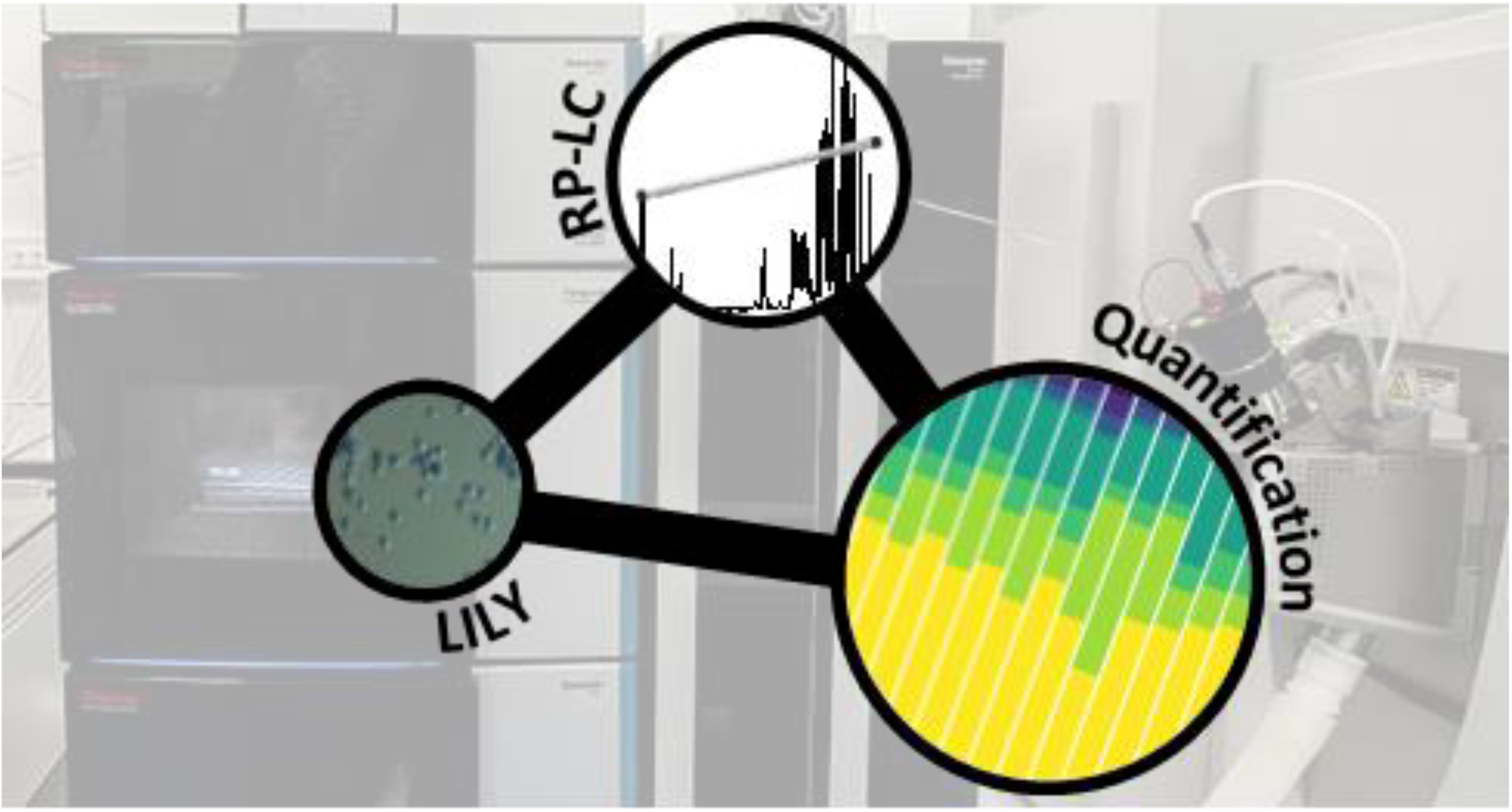

## INTRODUCTION

Lipidomics strives to comprehensively quantify lipids in biological entities. However, up to date, validation of quantitative omics-type of analysis remains a challenge.^1,2^ Proper standardization strategies regarding different analytical platforms and their compliance to guidelines are currently under debate. Internal standardization has been propagated as the method of choice for absolute quantification and the selection criteria of internal standards (ISTD) in lipidomics are well defined.^3^ Ideally, the lipid selected as ISTD is added as early as possible to the sample in the analytical process. In the best case it is the isotopically labeled analogue of the investigated lipid, otherwise at least the prerequisites of co-ionization and structural similarity have to be given. Different calibration approaches have been developed minimizing the number of required ISTDs: (1) Class-specific ISTD at a single concentration serve for the quantification of an entire lipid class,^4–10^ or (2) a class-specific ISTD is used at several concentration levels to obtain an internal calibration curve for the whole lipid class.^11–14^ Both calibration approaches are widely accepted for direct infusion mass spectrometry as all standards and analytes co-ionize. On top of that, the streamlined class-specific internal standardization is also applied in combination with chromatographic techniques separating lipids according to their head group chemistry (hydrophilic interaction liquid chromatography HILC, normal phase liquid chromatography NP-LC, supercritical fluid chromatography SFC), in which standards and analytes of the same lipid class co-elute and thus co-ionize.^15^ The commonly used ISTD panels^15^ comprise non endogenous lipid species with shorter fatty acyl chain(s) (e.g., 12:0 or 14:0), odd carbon number fatty acyl chain (e.g., 17:0, 17:1, 19:0) or endogenous deuterated analogues. Nowadays, dedicated one-lipid-per-class mixtures with optimized concentration levels for several sample types are available. As ionization efficiency is known to vary within one lipid class to some extent (the extent of this variation is depending on the lipid class)^16,17^ application of response factors compensating for these variations was proposed for neutral lipids.^18–23^ However, in most omics-type lipid analysis these corrections have been neglected.^24^ Targeted quantification of selected lipids based on external calibration with internal standardization by an (isotopically labeled) analogue is the method of highest metrological order.^25–29^ Only this approach complies with the FDA guideline for Bioanalytical Method Validation^30^ commonly used in clinics and biomarker evaluation. In fact, recent clinical large-scale studies have resorted to targeted analysis of a small panel of lipids.^31^

Nowadays, reversed phase-liquid chromatography mass spectrometry (RP-LC-MS) is the most widely used analytical lipidomics technique as revealed by a recent survey^1^ among expert laboratories and a comprehensive literature review.^24^ It is the method of choice when aiming at in-depth characterization of lipidomes. Reversed phase chromatography provides efficient matrix separation^32^ together with excellent chromatographic selectivity and retentivity for lipids, resulting in high sensitivity and dynamic range when combined to MS detection. While the method is unrivalled in terms of lipid separation and thus identification (in combination with high resolution MS (HRMS)), proper standardization approaches imply the use of multiple standards per lipid class. Lipid classes and potentially isomers (compounds with the same elemental composition)^33^ are separated according to their fatty acyl chain chemistry, as determined by chain length, the number of double bonds and their position. Consequently, quantification accuracy is compromised when only few standards per lipid class are implemented. ^3,15,32,34,35^

In this work we propose a RP-LC-HRMS workflow enlarging the number of standards in a cost saving manner, thereby improving the quantification capability of the most powerful lipidomics technique. More specifically, the presented standardization strategy is based on a lipid library provided by ^13^C fully labelled biomass, coined as lipidome isotope labeling of yeast (LILY).^25^ 250 uniformly ^13^C labeled lipid species from 19 lipid classes^25^ have been obtained after fermentation of the yeast strain *Pichia pastoris* (Guillierm.) Phaff 1956 (*Komagataella phaffii* Kurtzman).^36^ More recently, the number of identified lipids has been further increased to 405 lipid species applying a novel preparative-SFC workflow for lipid class fractionation^37^. An overlap of >100 lipid species with human plasma has been shown.^27^ While isotopically enriched biomass is frequently used in metabolomics,^38,39^ its value is less accepted in lipidomics. As quantitative information is lacking for LILY, the use as ISTD was restricted to internal standardization in combination with external calibrations. Accurate quantification based on the principles of isotope dilution has been enabled only by spiking external standards and samples with the same amount of LILY. The number of accurately quantified lipid species has been limited to the number of external calibrants.^26,27^ In order to overcome this limitation, a workflow has been designed to quantify LILY on an day-to-day routine prior to RP-LC-HRMS analysis of samples. This strategy enables to implement different calibration strategies within one lipidomics workflow including internal standardization without external standardization. Moreover, the recalibration of LILY coped with the fact, that comprehensive lipidome stability and storage conditions are still ill-defined for such cost-effective materials. It is well known, that certain lipid species are prone to oxidation and degradation making a frequent recalibration a prerequisite.^40–42^

Human plasma lipidomics serves as prime example to show the validity of the presented workflow as it represents the most frequently analyzed sample matrix in the field.^1,24^ The presented validation capitalizes on healthy donor samples ^43–47^ and reference materials^46^ with published quantitative lipidomes. However, a certified reference material ensuring traceability is still lacking in the field.

## MATERIALS AND METHODS

### Material

Human plasma samples were purchased from Innovative Research (Novi, Michigan). Standard reference material (SRM) 1950 from National Institute of Standards and Technology (NIST, Gaithersburg, USA) was used. Reference standards (endogenous compounds for external calibration) and SPLASH^®^ LIPIDOMIX^®^ Mass Spec Standard were obtained from Avanti (Alabaster, USA) and Merck KGaA (Darmstadt, Germany).

The internal standard LILY was obtained according to the procedure described in Neubauer *et al*^48^. and Schoeny *et al.*.^37^

### Method

Two different methods (FI and RP-LC) were applied in the same analytical sequence as enabled by a 6-port valve controlled via MS-software (see figure S1). A detailed description of extraction, analysis and data processing can be found in the extended material and methods section of the Supporting Information.

Briefly, a Vanquish Horizon HPLC and a high field Q Exactive HF™ quadrupole-Orbitrap mass spectrometer (both Thermo Fisher Scientific) were used. For RP chromatography of lipids, an Acquity HSS T3 (2.1 mm x 150 mm, 1.8 μm, Waters) with a VanGuard Pre-column (2.1 ~ 5 mm, 100 Å, 1.8 μm) were applied. The column temperature was set to 40°C and the flow rate to 250 μL min^−1^. Acetonitrile (ACN)/H_2_O (3:2, v/v) was used as solvent A and isopropanol (IPA)/ACN (9:1, v/v) as solvent B, both containing 0.1% formic acid and 10 mM ammonium formate. A gradient of 23 min was applied. MS1 acquisition was used for quantification. The injection volume of 2 μL was selected and polarity switching was performed. For data dependent acquisition (DDA) the LC method was identical, but the injection volume was increased to 5 μL, positive mode and negative mode were acquired separately and only the pooled sample together with the extraction blank and a high concentrated external standard were analyzed. For FI, the column was by-passed via 6-port valve. 25 μL were injected in a 5 μL min^−1^ flow to get a constant signal for around 5 min. The eluents were kept constant at 50% A/50% B. Each FI measurement lasted 10 min including washing. Polarity switching was triggered after 2.5 min (afterwards 10 sec for equilibration). For each polarity, only MS1 spectra were acquired at the beginning before 200 data independent acquisition (DIA) scans alternated with a MS1 scan for quantification. MS1 RP-LC lipid data was processed by Skyline (version 20.1), data dependent acquisition (DDA) files by LipidSearch 4.2 from Thermo Scientific and FI data was evaluated with LipidXplorer (version 1.2.8). All final data processing was performed in R/ R studio.

## RESULTS AND DISCUSSION

RP-LC-HRMS is the gold standard for in-depth characterization of lipidomes.^1,24^ While the superiority of this method with regard to sensitivity and dynamic range is widely accepted, its quantification capability is currently under debate. It becomes increasingly clear that a high number of lipid standards is mandatory for accurate - omics type of quantification by RP-LC based methods.^3,15,32,34,35^ In this work, a FI/LC-HRMS workflow integrated isotopically labeled yeast (LILY) as internal standards (see Figure 1) with the aim of expanding the number of lipids amenable to absolute accurate quantification as defined by the FDA guideline.^30^ Several proof-of-concept studies showed the potential of LILY.^25–27^ However, so far the applied internal standardization strategies lacked the quantitative characterization of the ^13^C yeast lipidome. Internal standardization was accomplished by spiking samples and external standards with known amounts of LILY.

**Figure 1.**
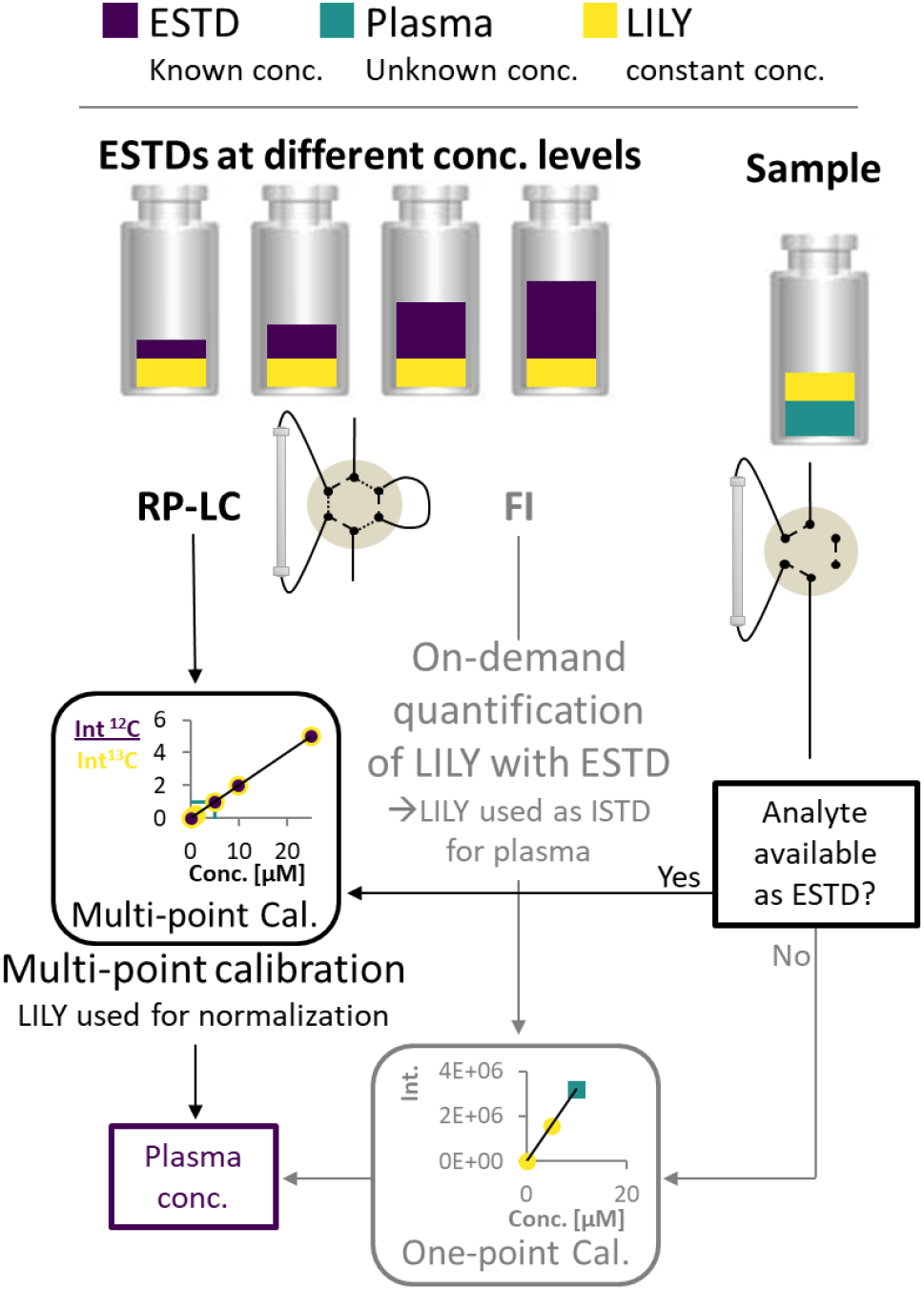
Overview of the combined quantification strategies. LILY (yellow) was added to the plasma sample (blue) and the ESTDs (purple) in the same concentration. ESTDs were measured via RP-LC-HRMS in MS1 mode and FI-MS to perform two different strategies: (1) a multi-point calibration internal standardized (normalized) with LILY lipids (RP data) and (2) an on-demand quantification of LILY lipids with the ESTD measured via FI-MS. Samples were only measured via RP-LC-HRMS. If the analyte is available as ESTD the first method can be used for quantification, if not the on-demand quantified LILY lipids were used as ISTD for one-point calibration.

### The combined FI/RP-LC-HRMS workflow for lipid quantification

In this work, a direct infusion analysis step preceded RP-LC-HRMS analysis and enabled quantitative characterization of LILY on a day to day basis by reverse isotope dilution. For this purpose flow injection and not the prevailing shotgun approach based on chip-solutions for nano-electrospray ionization (ESI)-MS was selected,^49^ given the possibility of automated switching to subsequent RP-LC based lipidomic analysis (see figure S1). An acquisition time of 5 min per sample was obtained by injecting 25 μL in a flow rate of 5 μL min^−1^ so that polarity switching and DIA MS/MS can be applied. As a prerequisite the UPLC system used in this work delivers highly precise flow rates in the flow regime from 1 μL min^−1^ to 5 mL min^−1^. Regarding sensitivity and precision, a comparable performance was observed for the optimized FI-HRMS approach and the chip-based infusion nano-ESI-MS^18^ (limits of detection (LOD) comparable to shotgun were found in this work; see also table S1).

Figure 1 depicts the principles of the FI/LC-HRMS workflow regarding standardization and measurement. The measurement sequence starts with direct infusion analysis of external standards spiked with LILY. Fully automated switching from FI analysis to RP-LC-HRMS is accomplished via a 6-port valve. A typical lipidomics RP-LC-HRMS run with 23 min was performed to analyze samples and external standards spiked with LILY lipids. Excellent retention time (RT) stability (see Figure S3A) supported lipid identification and quantification across samples. Despite the short chromatographic separation time, for some lipid species isomer separation was achieved for LPC, LPE, PC, PE and DG. Exemplarily, baseline separation of two lipid species (PC 18:1(9Z)/ 18:1(9Z) and PC 18:1(9E)/ 18:1(9E)) are shown in Figure S3B. However, quantification was based on the sum integral to simplify data evaluation. The streamlined workflow involved RP-LC-HRMS analysis of all samples and standards in full MS mode (MS1, mass resolution 120000, dynamic polarity switching) and of a pooled sample in data dependent MS/MS, for quantification and identification, respectively. A small loss of sensitivity due to the polarity switching routine in MS1 mode was accepted for the sake of saving analysis time. The measured calibration dilution series covered 4 orders of magnitude (low nM to low μM range). For data evaluation only the linear working range was considered.

Regarding the time distribution of the different tasks, almost a third of the time is spent on measuring samples, a quarter on quality control (QC) samples (including QC standards at four concentration levels, SRM 1950 and washing blanks), 20% on calibrants including replicates for limit of quantification (LOQ) determination (both on RP-LC (15%) and FI (5%)), 10% on DDA identification runs and 10% on system checks such as blank interferences and carry over tests. Overall, implementing FI-HRMS increased the RP-LC-HRMS workflow measurement time by less than 10% even when an extensive number of standards and different blanks were included (see figure S4 and Supporting Information Excel table/ Sequence). By tailoring calibration levels, the measurement time could be reduced even further.

As a key advance, the large panel of external and internal standards paved the way for different applicable calibration strategies expanding the number of lipid species amenable to absolute quantification. Moreover, an extensive dilution series of external calibrants allowed tailored calibration levels for lipid species evaluation, both in FI-HRMS and RP-LC-HRMS analysis. Thus, the method allowed accurate quantification without prior screening of linear range for each single analyte. Following stringent quality criteria, an automated selection of calibration strategy for each lipid species was implemented. Wherever applicable, species-specific standardization (denoted as level 1, according to the definition of LSI,^50^ see figure S5) was preferred over class-specific calibration (denoted as level 2 in case of RT matching, otherwise level 3). ISTD and ESTD were chosen in the following decreasing preference order: level of standard,^50^ number of hydroxy groups (necessary only for sphingolipids), number of double bonds and number of carbons in the fatty acyl chain. The quantification approach of highest metrological order is lipid species-specific standardization by external multi-point calibration with internal LILY standardization. Evidently, this calibration approach is limited to the selection of external standards and lipids present in LILY. Otherwise, level 2 (RT match was accepted for maximal +/− 0.5 min RT shift) external calibration using species-specific (level 1) LILY ISTD was applied followed by one-point calibration using the quantified level 1 LILY ISTD. If no species-specific calibration was available lipid class-specific RT matched standardization was applied. The remaining species were assessed using level 3 standards.

### Quantification of LILY by FI-HRMS

169 ^13^C fully labeled lipid species were identified on the lipid species level in the LILY extract using the proposed workflow. Spiking plasma samples with LILY upon extraction, increased the overall molar lipid amount of the sample by approximately 10%. Compared to naturally occurring isobaric overlaps in plasma between different lipid classes, adducts or isotopologues, the chance for isobaric overlaps between unlabeled and fully ^13^C labeled lipids is small. Theoretically, 48% of the plasma lipids have potential isobaric interferences with other lipids of the same sample but only 11% of them would be affected by labeled ^13^C LILY lipids if no separation is used.

67 LILY lipids species were accurately quantified by reverse isotope dilution on an FI-HRMS routine by using either level 1 or 2 type of standardization. Depending on the lipid class, MS quantification was based on the precursor, the head group fragment or the fatty acyl chain fragments in either positive or negative mode. Lipids from 8 classes (DG, TG, ST, PC, PE, PG, LPC, HexCer-see Supporting information for lipid class abbreviations) ranging at concentrations from 3 to 1500 nM were quantified (see Supporting Information excel table/LILY-lipids_FI). For all obtained LILY standard concentrations, stringent quality criteria were met in the FI-HRMS data. A detailed description of the quality criteria used for filtering is given in the Supporting Information. Overall, experimental uncertainty in QC samples, signal stability, linear dynamic range and LOQ were considered. Thus, out of the 169 LILY lipids, 67 compounds were potential one-point ISTDs with experimentally assessed concentrations. All other LILY lipids served as ISTD in external calibrations requiring no quantitative information to compensate for variation in sample preparation and instrument performance (see strategy 1 and 2 respectively in Supporting Information/absolute lipid quantification).

### Evaluation of different calibration strategies

Figure 2 shows concentration values for selected lipid species obtained by different calibration strategies in SRM 1950 with respect to the published consensus values.^46,51^ More specifically, the quantification of 4 lipid species, i.e. DG 34:1, TG 48:3, PE 36:2 and PC 36:2 is addressed considering external calibration with internal standardization and only internal standardization based on LILY lipids, both at different levels (level 1, 2, 3) and MS1 acquisition, respectively. The isomers of PC 36:2 were separated on the RP-LC. However, quantification was performed with the sum integral over all isomers. The importance of species-specific calibration or at least RT matching in class-specific calibration can be readily observed, emphasizing that a small number of standards is not practical in RP-LC analysis. Regardless if level 1 or level 2 calibration was applied all concentrations were within the 99% confidence interval as published for the SRM 1950 material. However, the international lipidomics interlaboratory comparison revealed rather broad distribution of measured concentration values for single lipid species^52^ (DG 34:1: 0-22 μmol L^−1^, TG 48:3: 1-10 μmol L^−1^, PE 36:2: 2-30 μmol L^−1^, PC 36:2: 75-350 μmol L^−1^) making it difficult to validate the different calibration strategies based on these consensus values only. Only certified reference material would allow an actual accuracy assessment.^44,53^

**Figure 2.**
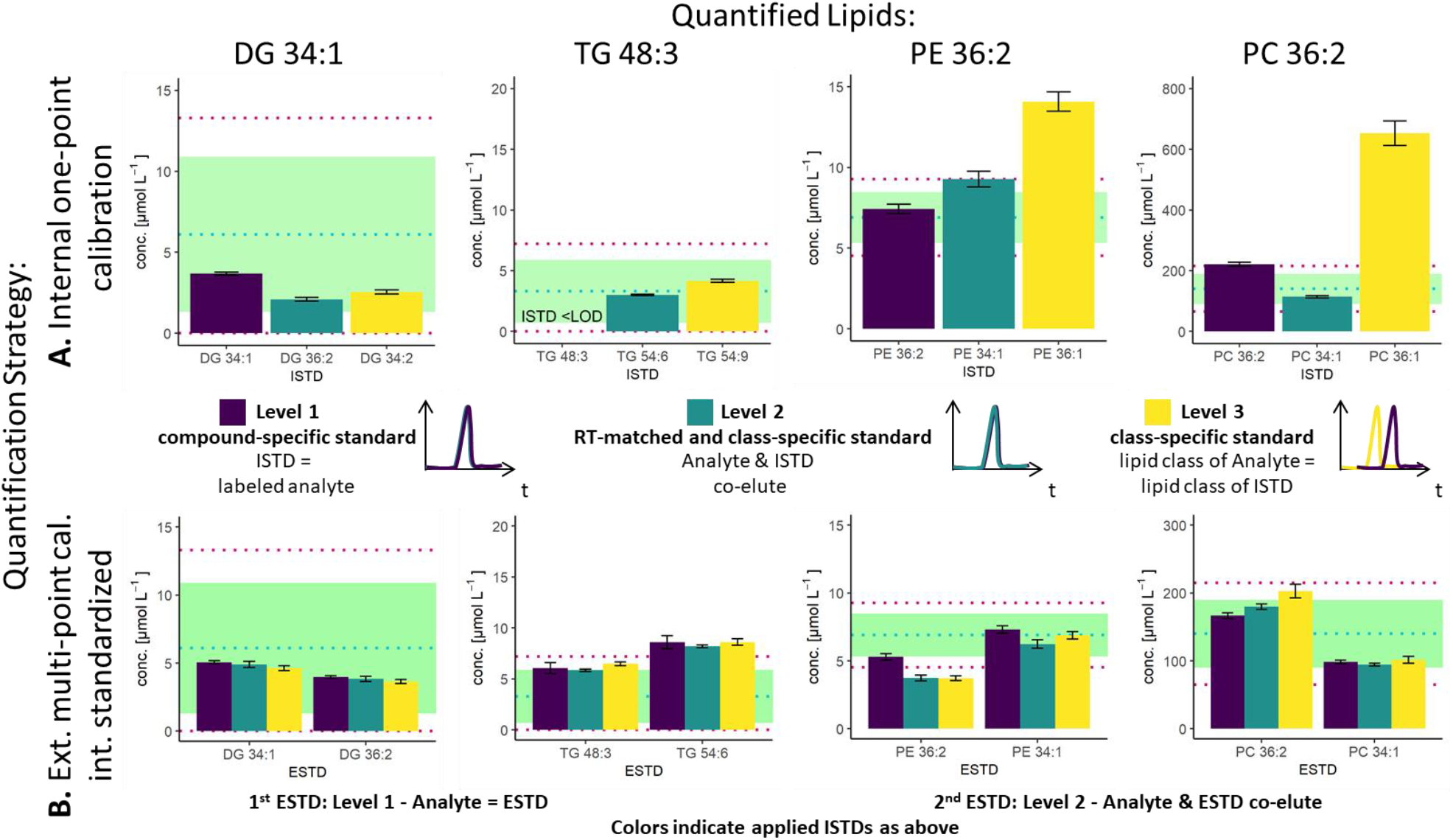
Influence of the selected ESTD and ISTD on (A) internal one-point calibration and (B) external multi-point calibration internal standardized. Different results were obtained with different levels of ESTD and ISTD. Exemplarily, the quantitative results of the lipids DG 34:1, TG 48:3, PE 36:2 and PC 36:2 in SRM 1950 are shown. The blue dotted line shows the median of the mean (MEDM), the light green region the 95% confident interval (CI) of the interlaboratory comparison and the red dotted line the 99% CI. (A) The x-axis and the color for the one-point calibration show the applied ISTD. (B) The x-axis for the multi-point calibration shows the applied ESTD (left: level 1, right: level 2) and the colors show the applied ISTD (purple: level 1, blue: level 2, yellow: level 3). The color code for the ISTD in B is equal to A.

As already mentioned, species-specific external calibration and internal standardization is the method of highest metrological order. The isotope dilution strategy ensures accuracy by compensating for losses during sample preparation and for variation of MS measurement, provided that (1) high purity external standards with certified concentrations are used and (2) equilibration of the LILY spike material and the sample is given. Considering typical experimental uncertainties of measured MS intensity ratios and sample preparation, the latter governed by extraction efficiencies and recoveries, typical total combined uncertainties of 4-7% were expected (see Figure 3). In fact, the experimentally observed uncertainties for biological replicates ranged at 2-7% when using level 1 ESTD and ISTD calibrations. When using level 2 external standardization with level 1 internal standardization, the ionization bias between the class-specific standard (ESTD) and investigated species contributes to the total combined uncertainty, while the contribution of sample preparation and MS detection was still minimized due to the species-specific ISTD. Introducing a correction factor - as estimate of the contribution of the ionization efficiency distribution within the lipid class- in the model equation of the uncertainty budgets results in estimated uncertainties of up to 30% for neutral lipids. However, for polar lipids this contribution was significantly lower resulting in calculated uncertainties ranging at 10% (ESTD level 2, ISTD level 1), in accordance to the widely accepted hypothesis that the ionization efficiency is determined through the head group.^3^ When using level 2 ESTD and level 2 ISTD, for polar lipids total combined uncertainties of 12-15% were estimated, while for neutral lipids again up to 35% were calculated. The combined uncertainty of species-specific internal standardization (one-point calibration) by LILY was mainly determined by the uncertainty of the LILY quantification based on FI-HRMS. Error propagation results in estimated combined uncertainties ranging at 14% (assuming that the measured ratios are within the dynamic range and level 1 standardization). Again, in the case of quantifying LILY via level 2 standardization, the uncertainty for neutral lipids increased to 33% as the differences in ionization efficiency cannot be overcome without correction. When performing species-specific ISTD (one-point calibration) with certified standards, 5-7% uncertainty have been estimated. It has to be mentioned that the estimated uncertainties of level 2 ISTD calibration is comparable regardless whether synthetic certified standards or cost saving LILY ISTD were used due to the fact that the correction factor for ionization bias is the major contribution to the total combined uncertainty. Consequently, the quantitative output of the workflow was ranked according to these considerations and labeled, accordingly. Quantification resorted to level 3 standardization only when level 1 /2 ESTD, ISTD were not available. Again, multi-point calibration was preferred over one-point calibration.

**Figure 3.**
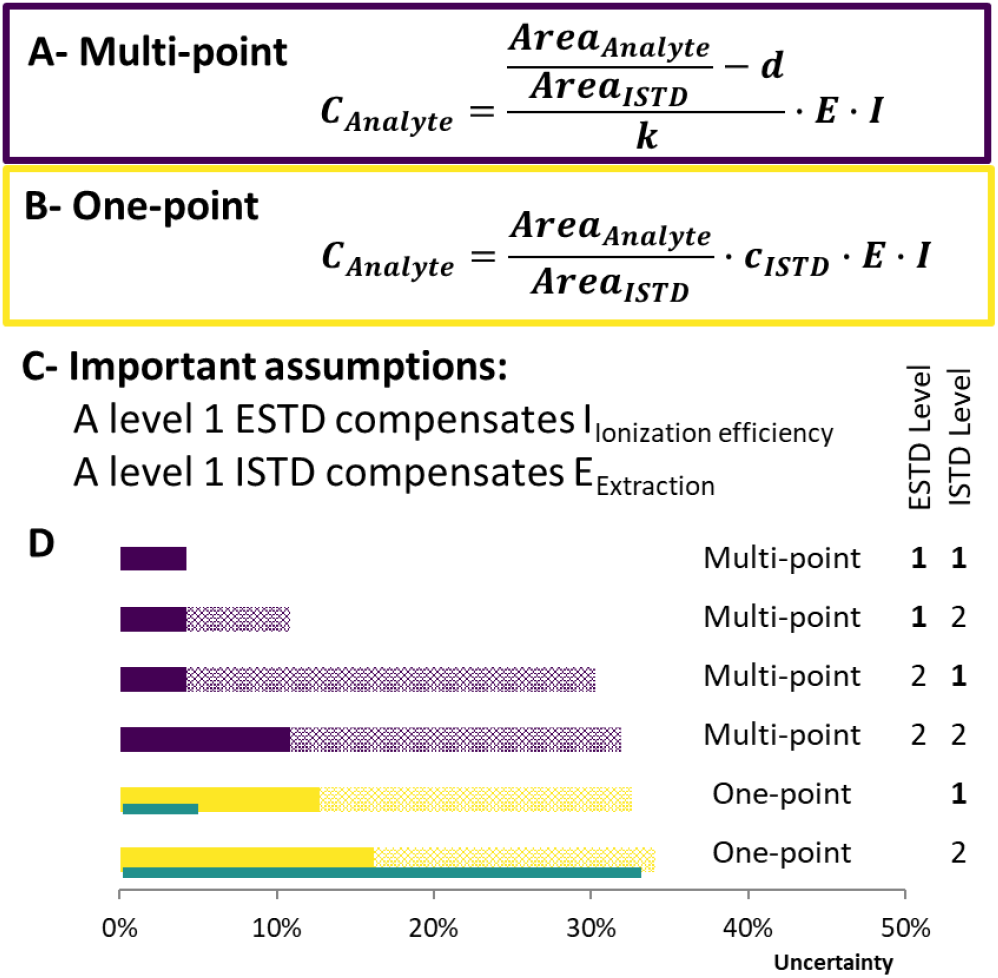
Uncertainty calculation of the applied methods. Concentration formula for (A) multi-point calibration and (B) one-point calibration. Area_Analyte_/Area_ISTD_: standard uncertainty of 3% in RP-LC, d-intercept, k-slope, E-Extraction factor E=1, associated uncertainty 10%,^54^ I-Ionization efficiency factor: I=1, associated uncertainty 30% (only for neutral lipids) **(C) Assumptions that have been made to estimate the uncertainty of each calibration. (D) ranking of the applied quantification strategies.** Purple bars indicate multi-point calibration, yellow one-point calibration. The blue bars next to one-point calibration show the uncertainty if synthetic certified standards (uncertainty of 2%) would have been used. Full color bars show the uncertainty for polar lipids whereas patterned stacked bars show the uncertainty that can occur for neutral lipid classes (e.g.: TG, CE, DG). One-point calibration considers LILY quantified upon level 2 standardization.

### Application to human plasma lipidomics

Although the yeast lipidome might be less complex, retention time windows matching to the respective plasma lipid classes were obtained (see Figure 4). Co-elution and so co-ionization was supported by the inverse retention order observed with respect to increased carbon number versus increased double bond number, also described in the equivalent carbon number (ECN) model.^55^ For example, TG 48:3 (ECN: 42) co-eluted with TG 50:4, TG 52:5, 54:6, etc. (all ECN: 42, exemplarily TG elution profile is shown in figure S3C, also true for DG, LPC, LPE, PC, PI). This phenomenon boosted the number of co-eluting LILY- and plasma lipid species. E.g. when analyzing the human plasma sample SRM1950, LILY provided a species-specific ISTD (level 1) for 26% out of the 357 quantified lipid species. For the great majority of lipid species (50 %), a class-specific and RT matched ISTD (level 2) was offered. Quantification of 8% of plasma lipids had to be based on a class-specific ISTD (level 3) only, while no ISTD was available for 16% (incl. SM, Hex2Cer and AcCa). In the latter cases external calibration had to be pursued. The selected ESTD panel enabled level 1 or 2 calibration for 50% out of the 16%. (see Figure 4).

**Figure 4.**
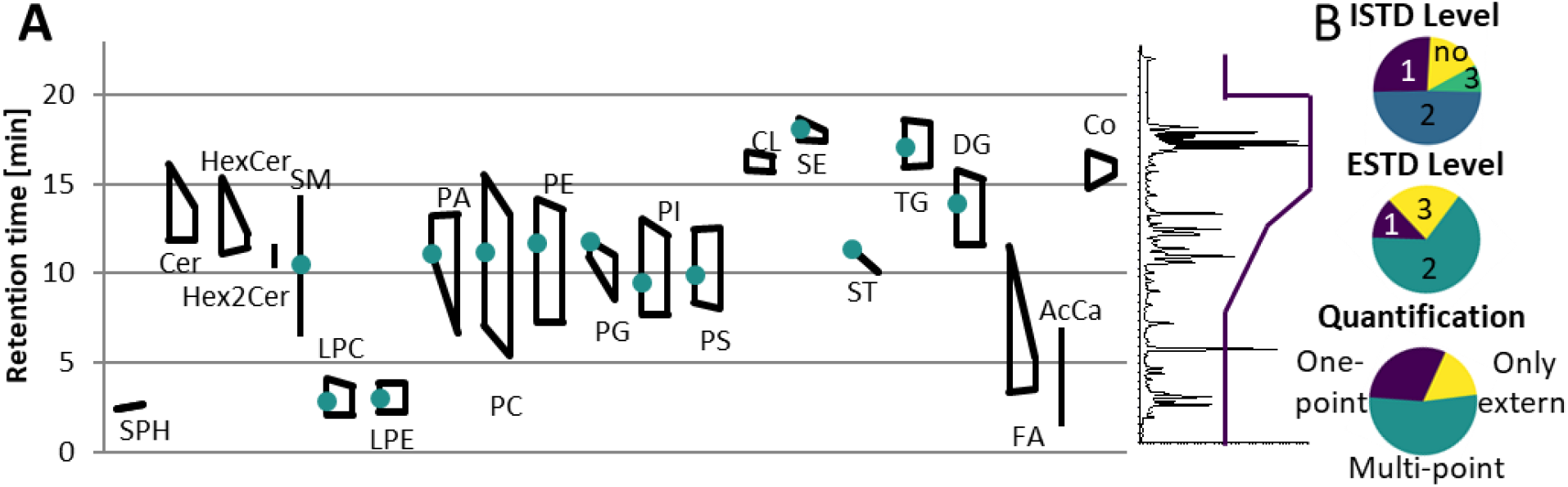
Application of LILY with RP-LC and its benefits (A) broad RT coverage: shows the RT coverage for different lipid classes, the left side of the quadrangle shows the elution range of human plasma lipids, the right side the elution range of LILY lipids, lipid classes shown as vertical line are only present in plasma, green dots show the elution time of the deuterated single-lipid-per-class mixture SPLASH^®^ LIPIDOMIX^®^ Mass Spec Standard. A total ion chromatogram and the gradient starting at 55% B and going up to 100% B is shown on the right side as visual help. **(B) Distribution of used ISTD, quantification strategies and ESTD. ISTD: 76% of analytes have RT matched ISTD**: shows the level (defined by the LSI)^50^ of ISTD. **Quantification strategies: 53% of analytes are quantified via multi-point calibration:** three different quantification strategies have been applied, multi-point: internal standardized multi-point calibration (according to FDA^30^), one-point: LILY is quantified in a multi-point calibration with ESTD first and analytes are quantified via one-point calibration with the calculated LILY conc. only extern: multi-point calibration without the use of ISTD **ESTD: 13 % of analytes are available as ESTD and 65% have co-ionized ESTD.**

**Figure 5.**
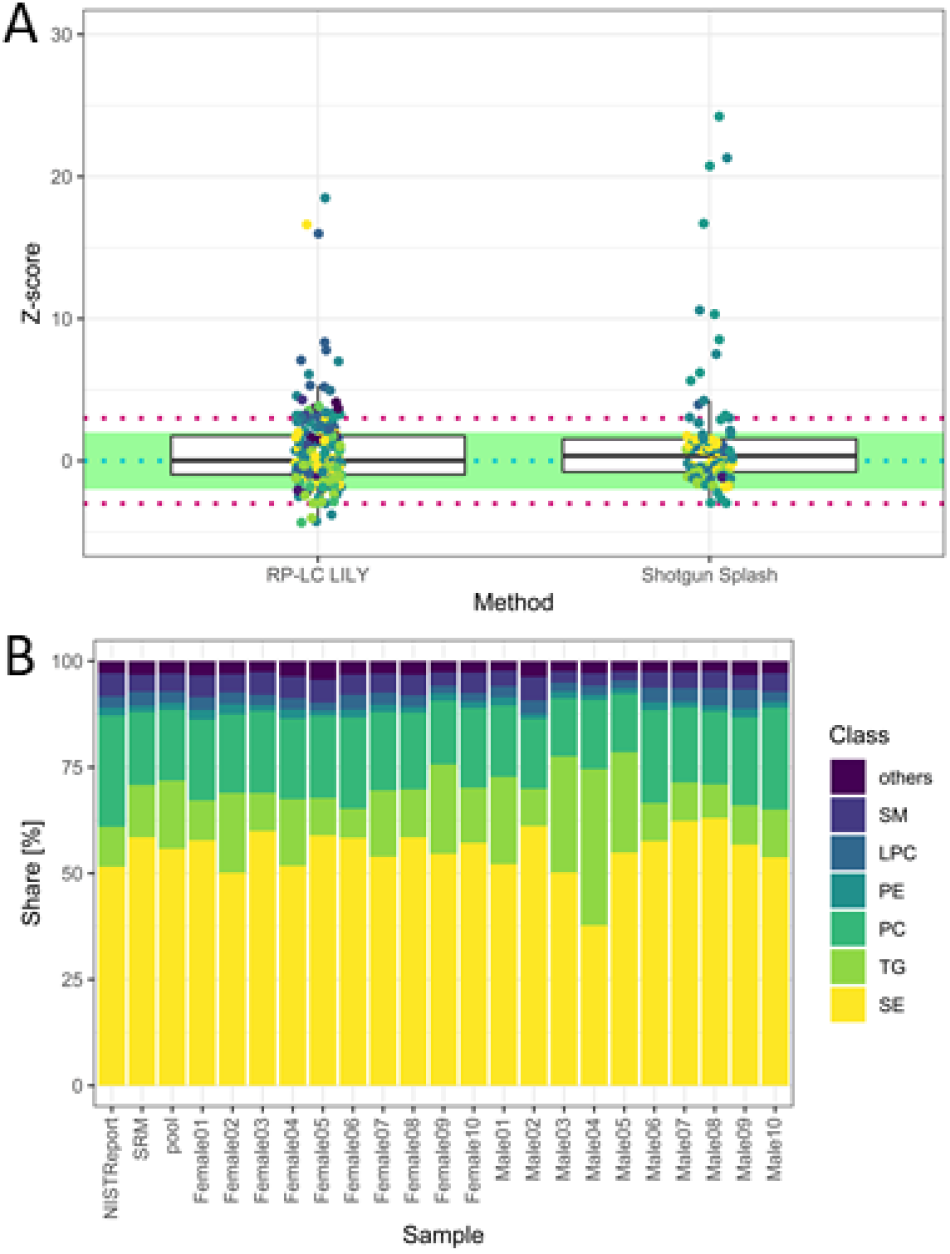
(A) Z-score distribution of the presented workflow (RP-LC LILY) covering 223 lipids in comparison with the state-of-the-art quantification using the SPLASH^®^ LIPIDOMIX^®^ in a shotgun nanoESI experiment covering 94 lipids. (coverage is determined by the NIST report). In figure S6 and table S2 a more detailed version can be found. Data for shotgun experiment taken from previous publication (Schoeny *et al.*^37^) **(B) Relative lipid class distribution of the major lipid classes in human plasma without cholesterol.**“Others” contains all lipid classes with a smaller contribution to the total lipid content then 2%. NIST report shows the distribution from the consensus values, whereas SRM is the analyzed SRM 1950 sample.

The FI/LC-HRMS lipidomics workflow was applied to the analysis to human plasma samples from 20 healthy donors striving to provide high lipidome coverage and accurate quantification. Lipid extraction of human plasma is well established.^15,56–59^ Spiking plasma with LILY increased the overall lipid amount of the sample by approximately 10%. Thus, the established Matyash protocol^60^ was adopted by reducing the plasma sample amount to a minimum of 10 μL, which still ensures homogeneity^40^ and increasing the sample versus solvent ratio to 1:325. By automatizing the workflow and streamlining the data evaluation, manual curation of data was reduced to a minimum (for more details see Supporting Information). The reference material for human plasma SRM 1950-providing consensus values for 339 lipid species based on an international interlaboratory comparison-was used for method validation and data filtering.^46^

Three independent methods (QC accuracy and precision tests, recovery test considering standard addition of non-endogenous standards and Z-score calculation of the SRM 1950) were implemented as quality control measures of the workflow next to the filter criteria described in the extended methods part in the Supporting Information. In total, quantitative values for 357 lipid species were obtained. 90 lipids from 7 lipid classes (LPC, LPE, PC, PE, PI, DG, TG) passed all quality filters. More specifically, accurate quantification was based on ESTD and ISTD panels passing the QC accuracy (accepting < 30% trueness bias) and precision tests (accepting < 30% relative standard deviation (RSD)), the recovery test with bias <30% for the respective lipid class and a calculated Z-score within the 99% CI.

The total quantitative output of the novel workflow was compared to the state-of-the-art shotgun lipidomics strategy^37^ with regard to the calculated Z-score based on SRM 1950 data. As can be readily observed in Figure 4A, the quantitative performance was comparable, after both data sets were filtered by strict criteria to avoid misidentifications including signal stability (RSD < 30%) and mass accuracy (< 3 ppm) and LOQ. However, the novel workflow benefits from increased sensitivity leading to a higher number of quantified lipids, an increased selectivity, the higher number of standards and the possibility to control accuracy by the mentioned tests.

The obtained relative lipid class distribution resembling the biological variance of the 20 healthy donors can be seen in Figure 4B.^44,61^ On average, state of the art lipidomics methods mentioned in literature reported 50–300 annotated lipid species when quantification was mentioned.^24^ In this work, 357 lipids out of 429 identified lipids were quantified in human plasma providing new tools for data validation and ranking quantitative values after a stringent metrological order. Additionally, new tools enabling data validation have been introduced, offering a new way of method control and improves the reliability of all quantitative results.

## CONCLUSIONS

Lipidome wide absolute quantification and validation was enabled via high resolution mass spectrometry coupled to reversed phase chromatography. As a prerequisite, implementation of a reversed isotope dilution step boosted the number of available lipid standards provided by fully labelled ^13^C yeast extracts in a cost-effective manner. Indeed, our experiments showed that the designed workflow provided a comparable analytical performance with regard to accuracy, uncertainty and number of absolute quantifications compared to established lipidomics platforms with the additional possibility to control accuracy via actual analyte QC accuracy and precision tests, recovery test considering standard addition of non-endogenous standards and Z-score calculation of the SRM 1950. Hence, all acknowledged benefits of RP-LC-HRMS based lipidomics, including isomer separation, superior sensitivity and selectivity are now amenable without compromising on the aspect of quantification. Thus, we think that the workflow increases the option of how to quantify in the field of lipid analysis offering an independent calibration strategy relying on different calibrants and standard resources, which is important regarding harmonization and standardization. Furthermore, this workflow can be used for any isotopically labeled biomass as ISTD enabling adaptation or self-production. Our study was restricted to quantification on a lipid species level based on MS1 measurements. In future, the quantification capability could be extended to the molecular species level (i.e. known fatty acyl chain composition) by integrating MS2 measurements and the LC-separated isomers in the workflow.

## SUPPORTING INFORMATION

PDF-Supporting information-Abbreviations, Extended Materials and Methods, Additional figures and tables

XLSX-quantitative results, quantification strategy comparison, figures of merit, quantified LILY lipids with FI-HRMS, the MS sequence list

ZIP-MFQL files for LipidXplorer

Raw data available on MetaboLights ^62^ www.ebi.ac.uk/metabolights/MTBLS1876

## ACKNOWLEDGMENTS

The Mass Spectrometry Center (MSC), Faculty of Chemistry, University of Vienna is acknowledged for providing mass spectrometric instrumentation. The authors thank all members of the environmental analysis (University of Vienna) group for continuous support.

## COMPLIANCE WITH ETHICAL STANDARDS

This work is supported by the University of Vienna, the Faculty of Chemistry, the Vienna Metabolomics Center (VIME; http://metabolomics.univie.ac.at/) and the research platform Chemistry Meets Microbiology. The project was funded from the aws PRIZE prototype funding by the Austrian BMWFW Federal Ministry. ISOtopic solutions supplied the yeast cells for internal standardization.

## Notes

https://www.ebi.ac.uk/metabolights/MTBLS1876

### Competing Interest Statement

The authors have declared no competing interest.

## REFERENCES

(1) Bowden, J. A.; Ulmer, C. Z.; Jones, C. M.; Koelmel, J. P.; Yost, R. A. NIST Lipidomics Workflow Questionnaire: An Assessment of Community-Wide Methodologies and Perspectives. Metabolomics 2018, 14 (53).

(2) Baker, P. R. Current State of Quantitation in Lipidomics Analysis https://avantilipids.com/discussions/current-state-of-quantitation-in-lipidomics-analysis (accessed Feb 12, 2020).

(3) Wang, M.; Wang, C.; Han, X. Selection of Internal Standards for Accurate Quantification of Complex Lipid Species in Biological Extracts by Electrospray Ionization Mass Spectrometry—What, How and Why? Mass Spectrom. Rev. 2017, 36 (6), 693–714. https://doi.org/DOI10.1002/mas.21492.

(4) Ejsing, C. S.; Sampaio, J. L.; Surendranath, V.; Duchoslav, E.; Ekroos, K.; Klemm, R. W.; Simons, K.; Shevchenko, A. Global Analysis of the Yeast Lipidome by Quantitative Shotgun Mass Spectrometry. Proc. Natl. Acad. Sci. U. S. A. 2009, 106 (7), 2136–2141. https://doi.org/10.1073/pnas.0811700106.

(5) Gallego, S. F.; Sprenger, R. R.; Neess, D.; Pauling, J. K.; Færgeman, N. J.; Ejsing, C. S. Quantitative Lipidomics Reveals Age-Dependent Perturbations of Whole-Body Lipid Metabolism in ACBP Deficient Mice. Biochim. Biophys. Acta 2017, 1862, 145–155. https://doi.org/10.1016/j.bbalip.2016.10.012.

(6) Gallego, S. F.; Højlund, K.; Ejsing, C. S. Easy, Fast, and Reproducible Quantification of Cholesterol and Other Lipids in Human Plasma by Combined High Resolution MSX and FTMS Analysis. J. Am. Soc. Mass Spectrom. 2018, 29 (1), 34–41. https://doi.org/10.1007/s13361-017-1829-2.

(7) Neubauer, C.; Sebastian, H.; Sessions, A. L.; Booth, I. R.; Bowen, B. P.; Dianne, K.; Dalleska, N. F. Towards Measuring Growth Rates of Pathogens during Infections by D2O‐labeling Lipidomics. Rapid Commun. Mass Spectrom. 2018, 32, 2129–2140. https://doi.org/10.1002/rcm.8288.

(8) Peng, B.; Ahrends, R. Adaptation of Skyline for Targeted Lipidomics. J. Proteome Res. 2016, 15, 291–301. https://doi.org/10.1021/acs.jproteome.5b00841.

(9) Shevchenko, A.; Simons, K. Lipidomics: Coming to Grips with Lipid Diversity. Nat Rev Mol Cell Biol 2010, 11 (8), 593–598. https://doi.org/10.1038/nrm2934.

(10) Surma, M. A.; Herzog, R.; Vasilj, A.; Klose, C.; Christinat, N.; Morin-Rivron, D.; Simons, K.; Masoodi, M.; Sampaio, J. L. An Automated Shotgun Lipidomics Platform for High Throughput, Comprehensive, and Quantitative Analysis of Blood Plasma Intact Lipids. Eur. J. Lipid Sci. Technol. 2015, 117, 1540–1549. https://doi.org/10.1002/ejlt.201500145.

(11) Gallego, S. F.; Hermansson, M.; Liebisch, G.; Hodson, L.; Ejsing, C. S. Total Fatty Acid Analysis of Human Blood Samples in One Minute by High-Resolution Mass Spectrometry. Biomolecules 2019, 9 (7), 1–16. https://doi.org/10.3390/biom9010007.

(12) Hammad, L. A.; Cooper, B. S.; Fisher, N. P.; Montooth, K. L.; Karty, J. A. Profiling and Quantification of Drosophila Melanogaster Lipids Using Liquid Chromatography/ Mass Spectrometry. Rapid Commun. Mass Spectrom. 2011, 25, 2959–2968. https://doi.org/10.1002/rcm.5187.

(13) Drotleff, B.; Hallschmid, M. ; Lämmerhofer, M. Quantification of Steroid Hormones in Plasma Using a Surrogate Calibrant Approach and UHPLC-ESI-QTOF-MS / MS with SWATH-Acquisition Combined with Untargeted Profiling. Anal. Chim. Acta 2018, 1022, 70–80. https://doi.org/10.1016/j.aca.2018.03.040.

(14) Drotleff, B.; Illison, J.; Schlotterbeck, J.; Lukowski, R. ; Lämmerhofer, M. Comprehensive Lipidomics of Mouse Plasma Using Class-Specific Surrogate Calibrants and SWATH Acquisition for Large-Scale Lipid Quantification in Untargeted Analysis. Anal. Chim. Acta 2019, 1086, 90–102. https://doi.org/10.1016/j.aca.2019.08.030.

(15) Holcapek, M.; Liebisch, G.; Ekroos, K. Lipidomic Analysis. Anal. Chem. 2018, 90, 4249–4257. https://doi.org/10.1021/acs.analchem.7b05395.

(16) Koivusalo, M.; Haimi, P.; Heikinheimo, L.; Kostiainen, R.; Somerharju, P. Quantitative Determination of Phospholipid Compositions by ESI-MS : Effects of Acyl Chain Length, Unsaturation, and Lipid Concentration on Instrument Response. J. Lipid Res. 2001, 42, 663–672.

(17) Hein, E.; Blank, L. M.; Heyland, J.; Baumbach, J. I.; Schmid, A.; Hayen, H. Glycerophospholipid Profiling by High-Performance Liquid Chromatography / Mass Spectrometry Using Exact Mass Measurements and Multi-Stage Mass Spectrometric Fragmentation Experiments in Parallel. Rapid Commun. Mass Spectrom. 2009, 23, 1636–1646. https://doi.org/10.1002/rcm.

(18) Höring, M.; Ejsing, C. S.; Hermansson, M.; Liebisch, G. Quantification of Cholesterol and Cholesteryl Ester by Direct Flow Injection High-Resolution Fourier Transform Mass Spectrometry Utilizing Species-Specific Response Factors. Anal. Chem. 2019, 91 (5), 3459–3466. https://doi.org/10.1021/acs.analchem.8b05013.

(19) Huynh, K.; Barlow, C. K.; Jayawardana, K. S.; Shaw, J. E.; Drew, B. G.; Meikle, P. J.; Huynh, K.; Barlow, C. K.; Jayawardana, K. S.; Weir, J. M.; Mellett, N. A.; Cinel, M.; Magliano, D. J.; Shaw, J. E.; Drew, B. G.; Meikle, P. J. High-Throughput Plasma Lipidomics : Detailed Mapping of the Associations with Cardiometabolic Risk Factors. Cell Chem. Biol. 2019, 26, 71–84. https://doi.org/10.1016/j.chembiol.2018.10.008.

(20) Weir, J. M.; Wong, G.; Barlow, C. K.; Greeve, M. A.; Kowalczyk, A.; Almasy, L.; Comuzzie, A. G.; Mahaney, M. C.; Jowett, J. B. M.; Shaw, J.; Curran, J. E.; Blangero, J.; Meikle, P. J. Plasma Lipid ProfiLing in a Large Population-Based Cohort. J. Lipid Res. 2013, 54, 2898–2908. https://doi.org/10.1194/jlr.P035808.

(21) Hofmann, T.; Schmidt, C. Instrument Response of Phosphatidylglycerol Lipids with Varying Fatty Acyl Chain Length in Nano-ESI Shotgun Experiments. Chem. Phys. Lipids 2019, 223 (104782). https://doi.org/10.1016/j.chemphyslip.2019.05.007.

(22) Schuhmann, K.; Moon, H.; Thomas, H.; Ackerman, J. M.; Groessl, M.; Wagner, N.; Kellmann, M.; Henry, I.; Nadler, A.; Shevchenko, A. Quantitative Fragmentation Model for Bottom-Up Shotgun Lipidomics. Anal. Chem. 2019, 91, 12085–12093. https://doi.org/10.1021/acs.analchem.9b03270.

(23) Tu, J.; Yin, Y.; Xu, M.; Wang, R.; Zhu, Z. J. Absolute Quantitative Lipidomics Reveals Lipidome-Wide Alterations in Aging Brain. Metabolomics 2018, 14 (5), 1–11. https://doi.org/10.1007/s11306-017-1304-x.

(24) Cajka, T.; Fiehn, O. Comprehensive Analysis of Lipids in Biological Systems by Liquid Chromatography-Mass Spectrometry. Trends Anal. Chem. 2014, 1 (61), 192–206. https://doi.org/10.1016/j.trac.2014.04.017.Cajka.

(25) Rampler, E.; Coman, C.; Hermann, G.; Sickmann, A.; Ahrends, R.; Koellensperger, G. LILY-Lipidome Isotope Labeling of Yeast: In Vivo Synthesis of ^13^C Labeled Reference Lipids for Quantification by Mass Spectrometry. Analyst 2017, 142, 1891–1899. https://doi.org/10.1039/C7AN00107J.

(26) Rampler, E.; Egger, D.; Schoeny, H.; Rusz, M.; Pacheco, M. P.; Marino, G.; Kasper, C.; Naegele, T.; Koellensperger, G. The Power of LC-MS Based Multiomics : Exploring Adipogenic Differentiation of Human Mesenchymal Stem/Stromal Cells. Molecules 2019, 24 (3615).

(27) Rampler, E.; Criscuolo, A.; Zeller, M.; El Abiead, Y.; Schoeny, H.; Hermann, G.; Sokol, E.; Cook, K.; Peake, D. A.; Delanghe, B.; Koellensperger, G. A Novel Lipidomics Workflow for Improved Human Plasma Identification and Quantification Using RPLC-MSn Methods and Isotope Dilution Strategies. Anal. Chem. 2018, 90 (11), 6494–6501. https://doi.org/10.1021/acs.analchem.7b05382.

(28) Ejsing, C. S.; Bilgin, M.; Fabregat, A. Quantitative Profiling of Long-Chain Bases by Mass Tagging and Parallel Reaction Monitoring. PLoS One 2015, 10 (12), 1–17. https://doi.org/10.1371/journal.pone.0144817.

(29) Demirkan, A.; Isaacs, A.; Ugocsai, P.; Liebisch, G.; Struchalin, M.; Rudan, I.; Wilson, J. F.; Pramstaller, P. P.; Gyllensten, U.; Campbell, H.; Schmitz, G.; Oostra, B. A.; Van Duijn, C. M. Plasma Phosphatidylcholine and Sphingomyelin Concentrations Are Associated with Depression and Anxiety Symptoms in a Dutch Family-Based Lipidomics Study. J. Pyschiatric Res. 2013, 47, 357–362. https://doi.org/10.1016/j.jpsychires.2012.11.001.

(30) Food and Drug Administration. Bioanalytical Method Validation Guidance for Industry; 2018.

(31) Kurz, J.; Parnham, M. J.; Geisslinger, G.; Schiffmann, S. Ceramides as Novel Disease Biomarkers. Trends Mol. Med. 2019, 25 (1), 20–32.

(32) Lange, M.; Ni, Z.; Criscuolo, A.; Fedorova, M. Liquid Chromatography Techniques in Lipidomics Research. Chromatographia 2019, 82, 77–100. https://doi.org/10.1007/s10337-018-3656-4.

(33) Wilkinson, A.; McNaught, A. D. IUPAC. Compendium of Chemical Terminology (the “Gold Book”), 2nd ed.; Blackwell Scientific Publications: Oxford, 1997. https://doi.org/doi.org/10.1351/goldbook.

(34) Hyötyläinen, T.; Ahonen, L.; Pöhö, P.; Oresic, M. Lipidomics in Biomedical Research-Practical Considerations. BBA - Mol. Cell Biol. Lipids 2017, 1862, 800–803. https://doi.org/10.1016/j.bbalip.2017.04.002.

(35) Burla, B.; Arita, M.; Arita, M.; Bendt, A. K.; Cazenave-gassiot, A.; Dennis, E. A.; Ekroos, K.; Han, X.; Ikeda, K.; Liebisch, G.; Lin, M. K.; Loh, T. P.; Meikle, P. J.; Oreši, M.; Quehenberger, O.; Shevchenko, A.; Torta, F.; Wakelam, M. J. O.; Wheelock, C. E.; Wenk, M. R. MS-Based Lipidomics of Human Blood Plasma: A Community-Initiated Position Paper to Develop Accepted Guidelines. J. Lipid Res. 2018, 59, 2001–2017. https://doi.org/10.1194/jlr.S087163.

(36) Kurtzman, C. P. Biotechnological Strains of Komagataella (Pichia) Pastoris Are Komagataella Phaffii as Determined from Multigene Sequence Analysis. J Ind Microbiol Biotechnol 2009, 36, 1435–1438. https://doi.org/10.1007/s10295-009-0638-4.

(37) Schoeny, H.; Rampler, E.; Hermann, G.; Grienke, U.; Rollinger, J. M.; Koellensperger, G. Preparative Supercritical Fluid Chromatography for Lipid Class Fractionation — a Novel Strategy in High-Resolution Mass Spectrometry Based Lipidomics. Anal. Bioanal. Chem. 2020, 412, 2365–2374.

(38) Schwaiger, M.; Schoeny, H.; Abiead, Y. El; Hermann, G.; Rampler, E.; Koellensperger, G. Merging Metabolomics and Lipidomics into One Analytical Run. Analyst 2019, 144, 220–229. https://doi.org/10.1039/c8an01219a.

(39) Hermann, G.; Schwaiger, M.; Volejnik, P.; Koellensperger, G. ^13^C-Labelled Yeast as Internal Standard for LC –MS / MS and LC High Resolution MS Based Amino Acid Quantification in Human Plasma. J. Pharm. Biomed. Anal. 2018, 155, 329–334. https://doi.org/10.1016/j.jpba.2018.03.050.

(40) Heiskanen, L. A.; Suoniemi, M.; Ta, H. X.; Tarasov, K.; Ekroos, K. Long-Term Performance and Stability of Molecular Shotgun Lipidomic Analysis of Human Plasma Samples. Anal. Chem. 2013, 85, 8757–8763. https://doi.org/10.1021/ac401857a.

(41) Yang, W.; Chen, Y.; Xi, C.; Zhang, R.; Song, Y.; Zhan, Q.; Bi, X.; Abliz, Z. Liquid Chromatography − Tandem Mass Spectrometry-Based Plasma Metabonomics Delineate the E Ff Ect of Metabolites ’ Stability on Reliability of Potential Biomarkers. Anal. Chem. 2013, 85, 2606–2610. https://doi.org/10.1021/ac303576b.

(42) Haid, M.; Muschet, C.; Wahl, S.; Prehn, C.; Adamski, J. Long-Term Stability of Human Plasma Metabolites during Storage at − 80 ° C. J. Proteome Res. 2018, 17, 203–211. https://doi.org/10.1021/acs.jproteome.7b00518.

(43) Pietzner, M.; Kaul, A.; Henning, A.; Kastenmüller, G.; Artati, A.; Lerch, M. M.; Adamski, J.; Nauck, M.; Friedrich, N. Comprehensive Metabolic Profiling of Chronic Low-Grade Inflammation among Generally Healthy Individuals. BMC Med. 2017, 15 (210), 1–12. https://doi.org/10.1186/s12916-017-0974-6.

(44) Triebl, A.; Burla, B.; Selvalatchmanan, J.; Oh, J.; Tan, S. H.; Chan, M. Y.; Mellet, N. A.; Meikle, P. J.; Torta, F.; Wenk, M. R. Shared Reference Materials Harmonize Lipidomics across MS-Based Detection Platforms and Laboratories. J. Lip 2020, 61, 105–115. https://doi.org/10.1194/jlr.D119000393.

(45) Cajka, T.; Smilowitz, J. T.; Fiehn, O. Validating Quantitative Untargeted Lipidomics Across Nine Liquid Chromatography–High-Resolution Mass Spectrometry Platforms. Anal. Chem. 2017, 89, 12360–12368. https://doi.org/10.1021/acs.analchem.7b03404.

(46) Bowden, J. A.; Heckert, A.; Ulmer, C. Z.; Jones, C. M.; Koelmel, J. P.; Abdullah, L.; Ahonen, L.; Alnouti, Y.; Armando, A.; Asara, J. M.; Bamba, T.; Barr, J. R.; Bergquist, J.; Borchers, C. H.; Brandsma, J.; Breitkopf, S. B.; Cajka, T.; Cazenave-Gassiot, A.; Checa, A.; Cinel, M. A.; Colas, R. A.; Cremers, S.; Dennis, E. A.; Evans, J. E.; Fauland, A.; Fiehn, O.; Gardner, M. S.; Garrett, T. J.; Gotlinger, K. H.; Han, J.; Huang, Y.; Huipeng Neo, A.; Hyötyläinen, T.; Izumi, Y.; Jiang, H.; Jiang, H.; Jiang, J.; Kachmann, M.; Kiyonami, R.; Klavins, K.; Klose, C.; Köfeler, H. C.; Kolmert, J.; Koal, T.; Koster, G.; Kuklenyik, Z.; Kurland, I. J.; Leadley, M.; Lin, K.; Maddipati, K. R.; Danielle, M.; Meikle, P. J.; Mellett, N. A.; Monnin, C.; Moseley, M. A.; Nandakumar, R.; Oresic, M.; Patterson, R.; Peake, D.; Pierce, J. S.; Post, M.; Postle, A. D.; Pugh, R.; Qiu, Y.; Quehenberger, O.; Ramrup, P.; Rees, J.; Rembiesa, B.; Reynaud, D.; Roth, M. R.; Sales, S.; Schuhmann, K.; Schwartzman, M. L.; Serhan, C. N.; Shevchenko, A.; Sommerville, S. E.; John-Williams, L. St.; Surma, M. A.; Takeda, H.; Thakare, R.; Thompson, J. W.; Torta, F.; Triebl, A.; Trötzmüller, M.; Ubhayasekera, K.; Vuckovic, D.; Weir, J. M.; Welti, R.; Wenk, M. R.; Wheelock, C. E.; Yao, L.; Yuan, M.; Zhao, X. H.; Zhou, S. Harmonizing Lipidomics: NIST Interlaboratory Comparison Exercise for Lipidomics Using Standard Reference Material 1950 –Metabolites in Frozen Human Plasma. J. Lipid Res. 2017, 58 (12), 2275–2288. https://doi.org/doi:10.1194/jlr.M079012.

(47) Thompson, J. W.; Adams, K. J.; Adamski, J.; Asad, Y.; Borts, D.; Bowden, J. A.; Byram, G.; Dang, V.; Dunn, W. B.; Fernandez, F.; Fiehn, O.; Gaul, D. A.; Fr, A.; Kalli, A.; Koal, T.; Koeniger, S.; Mandal, R.; Meier, F.; Naser, F. J.; Neil, D. O.; Pal, A.; Patti, G. J.; Pham-tuan, H.; Prehn, C.; Raynaud, F. I.; Shen, T.; Southam, A. D.; John-williams, L. S.; Sulek, K.; Vasilopoulou, C. G.; Viant, M.; Winder, C. L.; Wishart, D.; Zhang, L.; Zheng, J.; Moseley, M. A. International Ring Trial of a High Resolution Targeted Metabolomics and Lipidomics Platform for Serum and Plasma Analysis. Anal. Chem. 2019, 91, 14407–14416. https://doi.org/10.1021/acs.analchem.9b02908.

(48) Neubauer, S.; Haberhauer-Troyer, C.; Klavins, K.; Russmayer, H.; Steiger, M. G.; Gasser, B.; Sauer, M.; Mattanovich, D.; Hann, S.; Koellensperger, G. U^13^C Cell Extract of Pichia Pastoris - A Powerful Tool for Evaluation of Sample Preparation in Metabolomics. J. Sep. Sci. 2012, 35 (22), 3091–3105. https://doi.org/10.1002/jssc.201200447.

(49) Schuhmann, K.; Almeida, R.; Baumert, M.; Herzog, R.; Bornstein, S. R.; Shevchenko, A. Shotgun Lipidomics on a LTQ Orbitrap Mass Spectrometer by Successive Switching between Acquisition Polarity Modes. J. Mass Spectrom. 2012, 47 (1), 96–104. https://doi.org/10.1002/jms.2031.

(50) Lipidomics Standards Initiativ. Lipid Species Quantifciaton https://lipidomics-standards-initiative.org/guidelines/lipid-species-quantification (accessed Feb 12, 2020).

(51) Ulmer, C. Z.; Ragland, J. M.; Koelmel, J. P.; Heckert, A.; Jones, C. M.; Garrett, T.; Yost, R. A.; Bowden, A. LipidQC: Method Validation Tool for Visual Comparison to SRM 1950 Using NIST Interlaboratory Comparison Exercise Lipid Consensus Mean Estimate Values. Anal. Chem. 2017, 89, 13069–13073. https://doi.org/10.1021/acs.analchem.7b04042.

(52) Bowden, J. A.; Heckert, A.; Ulmer, C. Z.; Jones, C. M. Lipid Concentrations in Standard Reference Material (SRM) 1950: Results from an Interlaboratory Comparison Exercise for Lipidomics; 2017. https://doi.org/10.6028/NIST.IR.8185.

(53) Lange, M.; Fedorova, M. Evaluation of Lipid Quantification Accuracy Using HILIC and RPLC MS on the Example of NIST ^®^ SRM ^®^ 1950 Metabolites in Human Plasma. Anal. Bioanal. Chem. 2020, 412, 3573–3584.

(54) Patterson, R. E.; Ducrocq, A. J.; Mcdougall, D. J.; Garrett, T. J.; Yost, R. A. Comparison of Blood Plasma Sample Preparation Methods for Combined LC –MS Lipidomics and Metabolomics. J. Chromatogr. B jou 2015, 1002, 260–266.

(55) Lísa, M.; Holcapek, M. Triacylglycerols Profiling in Plant Oils Important in Food Industry, Dietetics and Cosmetics Using High-Performance Liquid Chromatography –Atmospheric Pressure Chemical Ionization Mass Spectrometry. J. Chromatogr. A 2008, 1199, 115–130. https://doi.org/10.1016/j.chroma.2008.05.037.

(56) Folch, J.; Lees, M. ; Sloane Stanley, G. H. A Simple Method for the Isolation and Purification of Total Lipides from Animal Tissues. J. Biol. Chem. 1957, 226, 497–509.

(57) Bligh, E. G.; Dyer, W. J. A Rapid Method of Total Lipid Extraction and Purification. Can. J. Biochem. Physiol. 1959, 37 (21), 911–917. https://doi.org/dx.doi.org/10,1139/cjm2014-0700.

(58) Ulmer, C. Z.; Jones, C. M.; Yost, R. A.; Garrett, T. J.; Bowden, J. A. Optimization of Folch, Bligh-Dyer, and Matyash Sample-to-Extraction Solvent Ratios for Human Plasma-Based Lipidomics Studies Candice. Anal. Chim. Acta 2018, 1037, 351–357. https://doi.org/10.1016/j.aca.2018.08.004.

(59) Han, X. Neurolipidomics: Challenges and Developments. Front Biosci. 2007, 12, 2601–2615.

(60) Matyash, V.; Liebisch, G.; Kurzchalia, T. V.; Shevchenko, A.; Schwudke, D. Lipid Extraction by Methyl-*Tert*-Butyl Ether for High-Throughput Lipidomics. J. Lipid Res. 2008, 49 (5), 1137–1146. https://doi.org/10.1194/jlr.D700041-JLR200.

(61) Quehenberger, O.; Armando, A. M.; Brown, A. H.; Milne, S. B.; Myers, D. S.; Merrill, A. H.; Bandyopadhyay, S.; Jones, K. N.; Kelly, S.; Shaner, R. L.; Sullards, C. M.; Wang, E.; Murphy, R. C.; Barkley, R. M.; Leiker, T. J.; Raetz, C. R. H.; Guan, Z.; Laird, G. M.; Six, D. a; Russell, D. W.; McDonald, J. G.; Subramaniam, S.; Fahy, E.; Dennis, E. a. Lipidomics Reveals a Remarkable Diversity of Lipids in Human Plasma. J. Lipid Res. 2010, 51 (11), 3299–3305. https://doi.org/10.1194/jlr.M009449.

(62) Haug, K.; Cochrane, K.; Nainala, V. C.; Williams, M.; Chang, J.; Jayaseelan, K. V.; Donovan, C. O. MetaboLights : A Resource Evolving in Response to the Needs of Its Scientific Community. Nucleic Acids Res. 2020, 48 (D1), 440–444. https://doi.org/10.1093/nar/gkz1019.

