## Supplementary material for "A combined flow injection/reversed phase chromatography – high resolution mass spectrometry workflow for accurate absolute lipid quantification with ^13^C- internal standards": Supporting Information.pdf

### 1. Abbreviations of lipid classes

| | | | |
|---|---|---|---|
| PL | phospholipid | NL | neutral lipid |
| PC | phosphatidylcholine | LPC | lysophosphatidylcholine |
| PE | phosphatidylethanolamine | LPE | lysophosphatidylethanolamine |
| PS | phosphatidylserine | LPS | lysophosphatidylserine |
| PG | phosphatidylglycerol | LPG | lysophosphatidylglycerol |
| PI | phosphatidylinositol | LPI | lysophosphatidylinositol |
| PA | phosphatidic acid | LPA | lysophosphatidic acid |
| CL | cardiolipin | ST | sterol |
| FA | fatty acid | DG | diglyceride |
| Cer | ceramide | TG | triglyceride |
| HexCer | hexosyl ceramide | MG | monoglyceride |
| Hex2Cer | dihexosyl ceramide | Co | coenzyme |
| SE | Sterol ester | CE | cholesterol ester |

### 2. Extended Materials and Methods

#### 2.1. Material

Human plasma samples were purchased from Innovative Research (Novi, Michigan). To prove the concept of the method 20 single plasma donors (10 male and 10 female) were analyzed. Standard reference material (SRM) 1950 from National Institute of Standards and Technology (NIST, Gaithersburg, USA) was used. Reference standards (endogenous compounds for external calibration) and SPLASH® LIPIDOMIX® Mass Spec Standard were obtained from Avanti (Alabaster, USA) and Merck KGaA (Darmstadt, Germany). An external standard mix produced from reference standards and used for quantification was produced and regularly controlled if lipids had been degraded by observing the ratio to SPLASH® LIPIDOMIX® lipids over time. The concentration was adopted if possible or the mixture was produced freshly. The internal standard lipidome isotope labeling of yeast (LILY) was obtained according to the procedure described in Neubauer *et al*[1]. and Schoeny *et al.*.[2] Other used chemicals were of LC-MS grade and ordered at Fisher Scientific (Vienna, Austria), VWR International (Vienna, Austria) or Sigma Aldrich (Vienna, Austria).

#### 2.2. Extraction

Extraction follows Matyash, *et al.*.[3] To a 10 µL plasma sample aliquot, which was placed at room temperature in a 5 mL microcentrifuge tube, 746 µL methanol (MeOH, cooled to -20°C), 4 µL SPLASH® LIPIDOMIX® Mass Spec Standard, 2 mL methyl tert-butyl ether (MTBE) (cooled to 4°C) and 0.5 mL of $^{13}C$ LILY extract (obtained from 1.6 $10^7$ cells) in MTBE were added. The mixture was shaken for 1h at room temperature. Phase separation was induced by adding 625 µL of MS-grade water. Upon 10 min of incubation, the sample was centrifuged at 1,000 rcf for 10 min. The upper (organic) phase was collected in a 5 mL microcentrifuge tube. The phase was dried under nitrogen and stored at -20°C if necessary. The dried lipids were dissolved in 200 µL IPA, centrifuged at 1000 rcf for 10 min and transferred into an insert in a HPLC vial. 10µL of each sample was pooled in a glass vial with insert to obtain a sample pool for lipid identification. An extraction blank was produced with the same protocol but 10 µL water instead of the plasma.

#### 2.3. Flow injection (FI) and reversed phase- liquid chromatography (RP-LC) high resolution mass spectrometry (HRMS) conditions

Two separate methods (flow injection (FI) and reversed phase- liquid chromatography (RP-LC)) can run in the same sequence as a fast switching was enabled by a 6-port valve controlled from the MS (see figure S1). Several parameters stayed constant during all measurements on the Vanquish Horizon and a high field Q Exactive HF™ quadrupole-Orbitrap mass spectrometer equipped with an electrospray both form Thermo Fisher Scientific. The injector needle was always washed with 75% isopropanol (IPA), 25% $H_2O$, 0.1% formic acid for 10 s prior and after each injection. This solution was also used for piston seal wash. Standards were measured in increasing concentration order, samples were randomized.

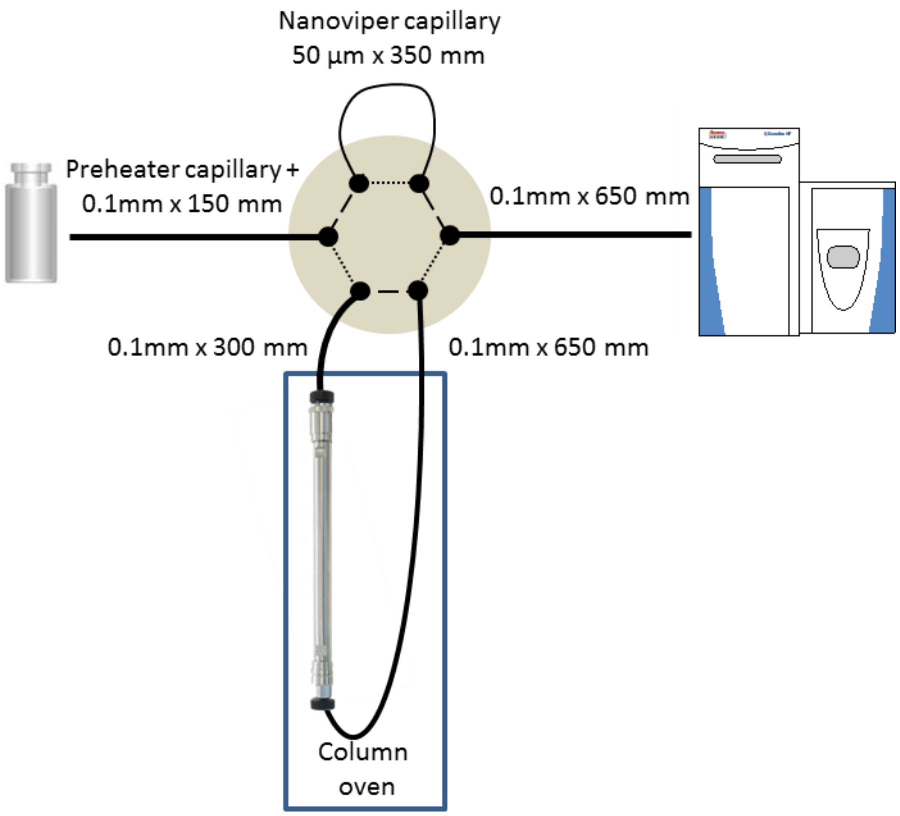

**Figure S1 6-port valve for the switch between RP-LC (dotted line) and DI (dashed line) with the dimensions of the used capillaries**

For RP chromatography of lipids, an Acquity HSS T3 (2.1 mm x 150 mm, 1.8 µm, Waters) with a VanGuard Pre-column (2.1 x 5 mm, 100 Å, 1.8 µm) was used. The column temperature was set to 40°C and the flow rate to 250 µL $min^{-1}$. Acetonitrile (ACN)/$H_2O$ (3:2, v/v) was used as solvent A and IPA/ACN (9:1, v/v) as solvent B, both containing 0.1% formic acid and 10 mM ammonium formate. The following gradient was applied: start at 55% B, 0−8.0 min ramp to 65% B, 8.0-13.0 min ramp to 85% B, 13.0-15.0 min ramp to 100% B, 15.0-20.0 min 100% B, 20.0 min fast switch to 55% B and equilibrated at the starting conditions for 3 min (20.0-23.0 min 55% B).

MS1 acquisition was used for quantification. The injection volume of 2 µL was selected and polarity switching was performed. The electrospray ionization (ESI) source parameters were the following: sheath gas 35, auxiliary gas 5, spray voltage 3.1 kV in both modes, capillary temperature 220 °C, S-Lens radio frequency (RF) level 30 and auxiliary gas heater 300 °C. Spectral data were acquired in profile mode. The full MS runs in positive and negative mode were acquired in the range of *m/z* 200 – 2000 at a resolution of 120,000, an automatic gain control (AGC) target at 1e6 and a maximum injection time (IT) of 200 ms.

For data dependent acquisition (DDA) the LC method was identical, but the injection volume was increased to 5 µL, positive mode and negative mode were acquired separately and only the pooled sample together with the extraction blank and a high concentrated external standard were analyzed. The MS parameters were the following: For both polarities, a Top8 method with normalized collision energy (NCE) of 25 (+)/28 (-) and an isolation window of *m/z* 1 was applied. In MS1 the range of *m/z* 200-2000 at a resolution of 120,000 was used with an AGC target at 1e6 and a maximum IT of 200 ms. The resolution in the MS2 was set to 30,000, the AGC target to 2e5 (minimum 8e3) and the max IT to 60 ms. The dynamic exclusion of triggered *m/z* was set to 15 s. An inclusion list was used for all possible lipids in human plasma. An exclusion list was generated by acquisition of a solvent blank run.

For FI, the column was by-passed via 6-port valve. 25 µL were injected in a 5 µL $min^{-1}$ flow to get a constant signal for around 5 min. The eluents were kept constant at 50% A/50% B. The flow starts at 100 µL $min^{-1}$ for 0.1 min, after a fast switch to 5 µL $min^{-1}$ this flow was kept until 5.5 min. The flow was switched to 250 µL $min^{-1}$ for flushing until 9 min at which it was switched back to 100 µL $min^{-1}$, which was kept until the end at 10 min.

Following tune settings were crucial for a stable flow: ionization voltage 3.5 kV (+)/-2.8 kV (-); sheath gas flow rate of 5; aux gas heater temperature 50°C; aux gas flow rate of 10; sweep gas flow rate of 4 capillary temperature 200°C; S-Lens RF level 50.

Each standard mixture was measured for 10 min and polarity switching was triggered after 2.5 min (afterwards 10 sec for equilibration). For each polarity, only MS1 spectra were acquired at the beginning before 200 data independent acquisition (DIA) scans alternated with a MS1 scan for quantification. In MS1, the resolution was set to 240,000, the AGC target to 1e6 and the maximum IT to 150 ms. For the DIA scans, a resolution of 60000 was applied and the AGC target and the max IT was set to 2e5 (+)/-5e5 (-) and 130 ms, respectively. NCE of 25 was used in positive mode and 28 in negative mode. The scan range was set to *m/z* 250-1200 in both modes.

#### 2.4. RP-LC-MS1 data pre-processing by Skyline

Quantitative RP-LC lipid data was processed by Skyline (version 20.1). Raw files were converted by MSConvert (Proteowizard) into mzML (centroided). A transition list was uploaded to Skyline in the molecule interface containing all possible analytes, external standards (ESTDs) and internal standards (ISTDs). Selected lipids were based on identification software outputs, elution order, main adduct abundance, polarity matching, the awareness of common misidentifications (see website of the Lipidomics Standard Initiative (LSI)).[4] All MS1 files were imported and peak boundaries were chosen manually in order to obtain accurate peak integration. The results were exported as csv-report (columns: File Name, Acquired Time, Precursor Ion Name, Molecule Formula, Isotope Label Type, Best Retention Time (RT), Total Area, Total Area Ratio). ISTDs with severe ion suppression were excluded from the RP-LC-HRMS data by comparing the signal in the spiked sample and the spiked extraction blank by setting a threshold.

#### 2.5. FI-MS data pre-processing by LipidXplorer

Data evaluation for identification and quantification of FI data was performed with LipidXplorer (version 1.2.8). Raw files were converted by MSConvert following the guidelines of the LipidXplorer wiki. The converted mzML files were imported via LipidXplorer into a Master Scan database using following settings: mass tolerance 5 ppm (MS1)/ 0.02 Da (MS2), min. occupation of 0, intensity threshold 0 (MS1)/ 0 (MS2), resolution 230000 (MS1)/ 60000 (MS2), resolution gradient -90 (MS1)/ -20 (MS2) in positive mode and -170 (MS1)/ -40 (MS2). The molecular fragmentation query language (MFQL) files used for identification can be found in the Supporting Information. The Reporting of all MFQL files needs to be equal for further R data evaluation. The following information was reported:

MASS (*m/z* value), CHEMSC (formula of ion), FORMULA (formula of neutral lipid species), ERROR (mass error between accurate and exact mass), CLASS (lipid class), NAME (shorthand notation), SPECIES (shorthand notation on fatty acyl chain level), ISOBARIC (occurrence of isobaric overlaps with other identified lipids), PRINTENS (intensity of precursor ion), depending on lipid class: Head intensity/fatty acid (FA) intensity, Headmass/FAmass, Headerror/FAerror. The data was exported as csv-file.

#### 2.6. RP-LC-HRMS lipid identification by LipidSearch

A fast lipid screening was performed with LipidSearch 4.2 from Thermo Scientific, by analyzing the DDA files only. Following filters were applied: RT tolerance 0.25 min, m-score threshold 2, ID quality filter A,B,C (D- only for free fatty acids and cardiolipins), calculate unassigned peak area TRUE and toprank filter TRUE. The identifications were curated manually following the criteria published elsewhere.[2] The data was exported as txt-file.

#### 2.7. Absolute lipid quantification

By automatizing the workflow and streamlining the data evaluation, manual curation of data was reduced to a minimum. All data processing was performed in R/ R studio. A brief summary is given here.

RP-LC-HRMS data: From the Skyline export, information about the file and the analyte were added from the file name or the formula, blank values were subtracted and the stability of area and RT of the quality control (QC) standards (10, 100, 1000 and 10000 nM) was calculated and filtered by certain thresholds (area relative standard deviation (RSD) < 30%, RT RSD < 3%). The area was also deisotoping Type 1 corrected[5] and compounds with a mass error above 3 ppm were removed. To control the RT and the lipid identification, LipidSearch data was integrated and mismatching lipids were reported for manual control. The lipid identification output of the utilized software packages, was curated considering elution order, main adduct abundance, polarity matching, and finally being aware of common misidentifications (see website of the LSI).[4]

FI-HRMS data: The csv export from LipidXplorer was transformed, information was added as with the Skyline data, average blank values for each analyte were calculated and subtracted, the intensity stability of the lipids in the QC standards (10, 100, 1000 and 10000 nM) were controlled (RSD < 30%), compounds with a mass error above 3 ppm were removed and deisotoping Type I[5] and Type II (adopted script published by Triebl *et al.*[6]) were performed. For data evaluation only the linear working range of the standard addition dilution series was considered and only LILY concentrations above the lower limit of the linear range (see Figure 2) were reported. We defined the lower limit of quantification (LOQ) and the upper one (ULOQ) according to FDA guideline recommendations[7] as it was deduced from the linear working range. Figure 2 also shows the automatized solution to determine the linear range by calculating the slope between all neighboring points and remove of all points at the ends until the single slopes are higher than 10% of the average slope. The limit of 10% was chosen according to visual inspection.

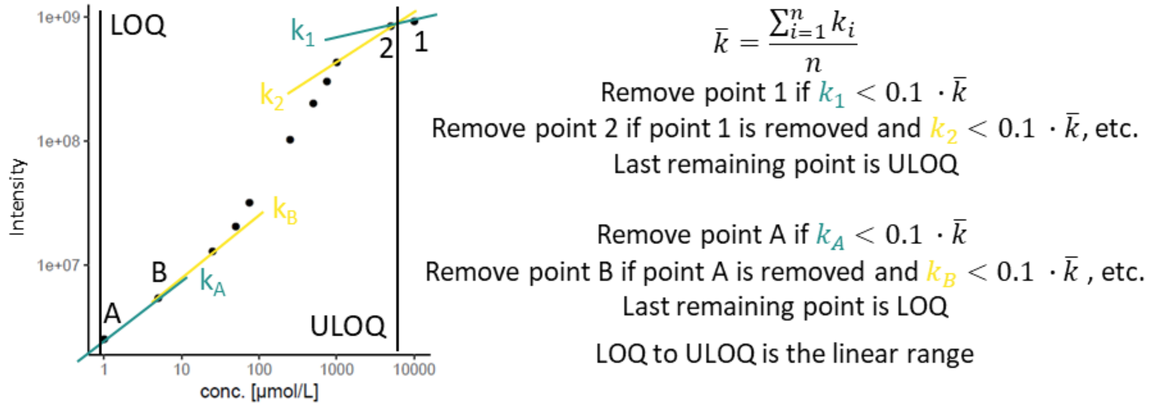

**Figure S2 Determination of the linear range and the LOQ/ULOQ in FI-HRMS measurements.**

For an easier nomenclature, the 3 levels definition of the LSI[4] (see figure S5) was used for ESTD and ISTD. Only the standards containing ESTD at a minimum of 6 concentration levels were used and LILY lipids were matched to the available ESTD based on the level of standard, number of hydroxy groups, number of double bonds and number of carbons in the fatty acyl chain. Always the closest 6 concentration levels were finally used for calibration. On top of that, for the final selection of LILY standards, only species with area RSD < 30% and RT RSD < 3% in the RP-LC-HRMS data as assessed in replicate QCs were accepted. The remaining quantified LILY lipids can be used as one-point calibrators for the plasma sample (second quantification strategy in the following paragraph).

**Three quantitative strategies were applied:**

##### (1) Multi-point calibration using the ESTDs measured via RP-LC-HRMS internal standardized with LILY lipids

Each LILY ISTD was matched to an ESTD of the same class based again on the level of standard, number of hydroxy groups, number of double bonds and number of carbons in the fatty acyl chain. The area ratios of ESTD/ISTD were calculated with the RP-LC-HRMS data only (one for each ISTD), and the linearity was determined as above. Area ratios between analyte and ISTD were calculated selecting the ISTD according to the previously described rules. Linear regression was calculated considering for analyte/ISTD ratio the six closest calibration points of the corresponding ESTD/ISTD (see equation 1). The LOQs of RP-LC-HRMS data were calculated by multiplying the standard deviation of low ESTDs (at conc. 1, 10 or 100 nM, n=5) with a factor of 10 and calculating a concentration based on the multi-point calibration. Concentrations below LOQ were discarded.

$$c_{Analyte} = \frac{\frac{Area_{Analyte}}{Area_{ISTD}} - d}{k} \tag{1}$$

##### (2) One-point calibration with the on-demand quantified LILY lipids as ISTD

Within this strategy, the via FI-HRMS quantified LILY ISTDs were applied as one-point calibrators. Again, the ISTD and the analytes were matched according to the rules above, concentration was calculated (see equation 2) and concentrations below LOQ were discarded.

$$c_{Analyte} = \frac{Area_{Analyte}}{Area_{ISTD}} \cdot c_{ISTD} \tag{2}$$

##### (3) Only external standardization based multi-point calibration using the ESTDs measured via RP-LC-HRMS (for lipid classes not present in LILY)

This strategy follows the same principle as strategy 1 but no ISTD were used. Therefore, the ESTD were directly matched with the analytes following again the rules above. The area of the six closest ESTDs were used to calculate the regression equation and the final lipid concentration of each analyte (see equation 3). LOQs were determined as strategy 1 and concentrations below LOQ were discarded.

$$c_{Analyte} = \frac{Area_{Analyte} - d}{k} \tag{3}$$

The different quantification strategies were calculated independently. Quantitative values were accepted provided that the recovery test of a non-endogenous standard was accurate for each class and the Z-score calculation of the SRM 1950 NIST report was within the 99% confident interval (CI) for each lipid species. For each analyte, concentration values based on different calibration strategies were obtained (see Supporting Information Excel table) and the final value was chosen based on the order following the uncertainty calculation.

### 3. Limit of detection (LOD) comparison

**Table S1 LOD comparison of shotgun methods (FI and chip based Nanomate robot system) and RP-LC in positive mode and during pos/neg switching**

| Class | Name | LOD [nmol $L^{-1}$] FI | Nano-mate | LC pos | LC posneg |
|---|---|---|---|---|---|
| DG | DG 34:1 | 5.0 | - | 6.3 | 3.5 |
| DG | DG 36:2 | 21 | - | 1.2 | 1.7 |
| LPC | LPC 16:0 | - | - | 1.4 | 17 |
| LPE | LPE 18:0 | - | - | 2.1 | 13 |
| PC | PC 32:0 | - | - | 17 | 56 |
| PC | PC 34:1 | 55 | - | 2.8 | 6.5 |
| PC | PC 34:2 | 64 | 12 | 0.6 | 2.2 |
| PC | PC 36:2 | - | - | 1.0 | 9.2 |
| PE | PE 34:1 | 95 | 60 | 2.0 | 21 |
| PE | PE 36:2 | 19 | - | 1.0 | 8.7 |
| TG | TG 48:3 | 22 | 13 | 2.0 | 1.0 |
| TG | TG 54:3 | 9.5 | - | - | 49 |
| TG | TG 54:6 | 25 | 25 | - | 16 |
| TG | TG 54:9 | 10 | 6.1 | 0.1 | 3.3 |
| TG | TG 60:3 | - | - | 0.4 | 22 |

#### 4. Chromatographic figures

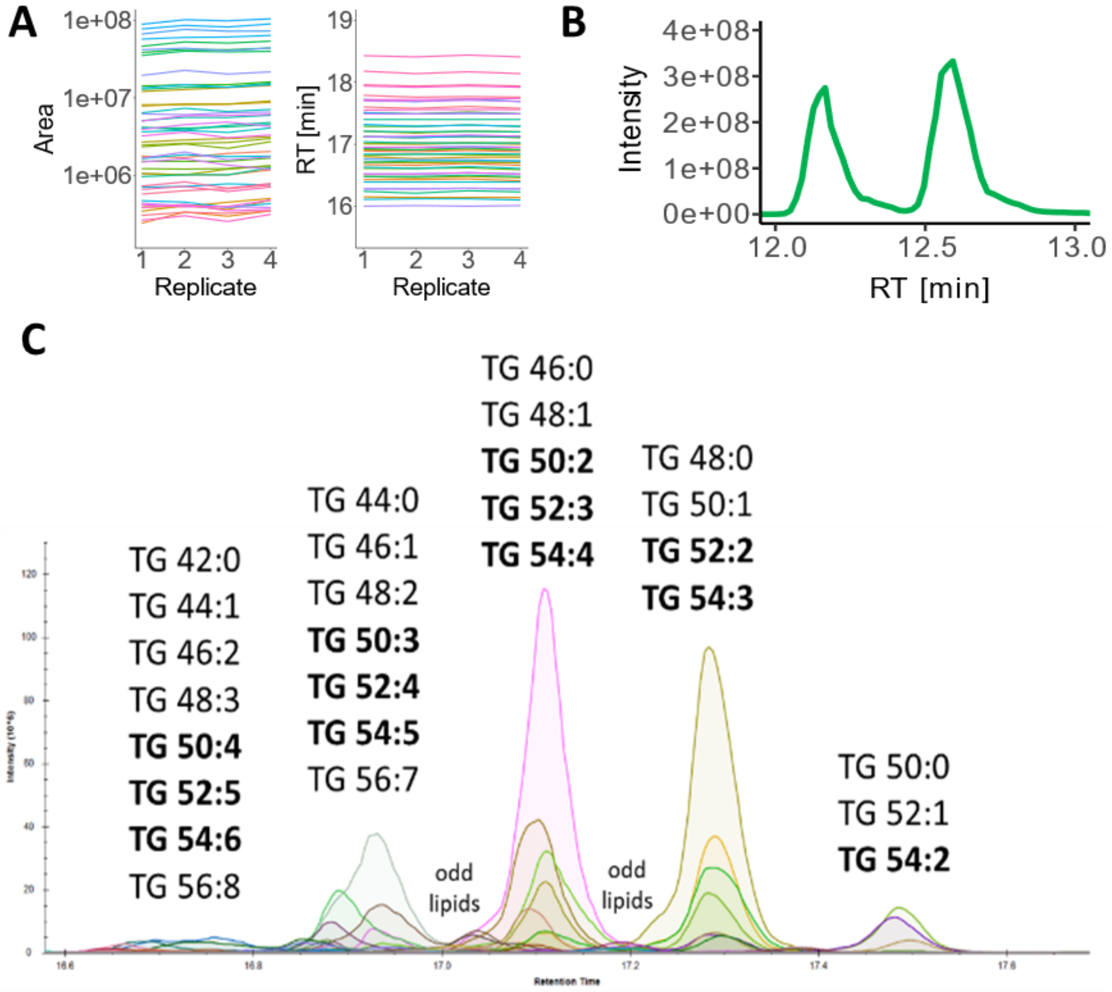

**Figure S3 (A) stable analysis:** RT and area stability over the run (3 days). **(B) Isomer separation:** baseline separation of PC 18:1(9Z)/ 18:1(9Z) and PC 18:1(9E)/ 18:1(9E). **(C) Retention time coverage of plasma lipids and LILY-ISTD.** shows the elution profile of TG plasma lipid species. Lipids, which occur also in yeast, are written in bold.

#### 5. Sequence overview

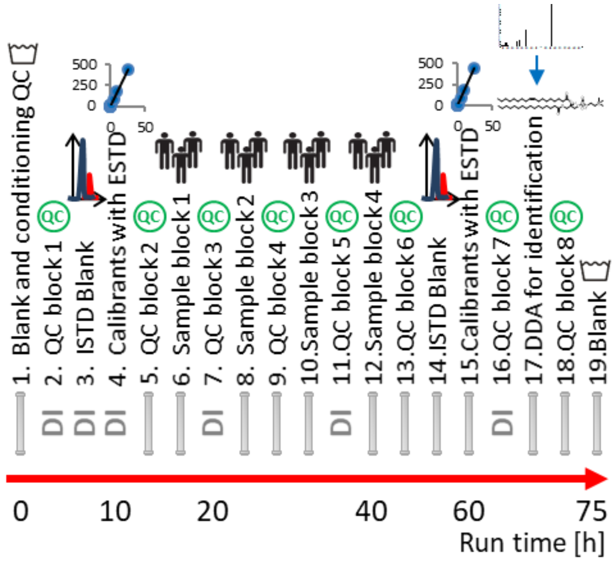

**Figure S4 Sequence overview.** Should give an overview of a suggested sequence for sample measurement. A detailed list can be found in the Supporting Information Excel file. Briefly, the column was washed and conditioned at the beginning followed by the calibrants measured via DI-MS, the samples divided in four blocks and the calibrants measured via RP-LC-MS. Between each block or after 10 injections, QC samples were measured alternately via DI or RP-LC. At the end, the pooled sample was measured with DDA for lipid identification and the column was washed with blank injections.

### 6. Levels of quantification

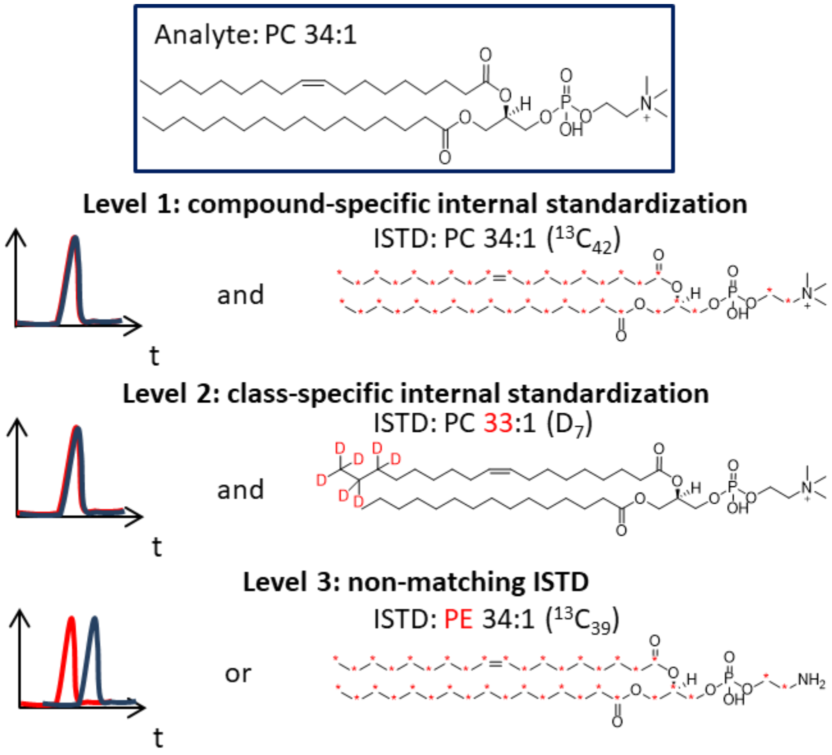

**Figure S5 Levels of Quantification as defined by the LSI**

### 7. SRM control

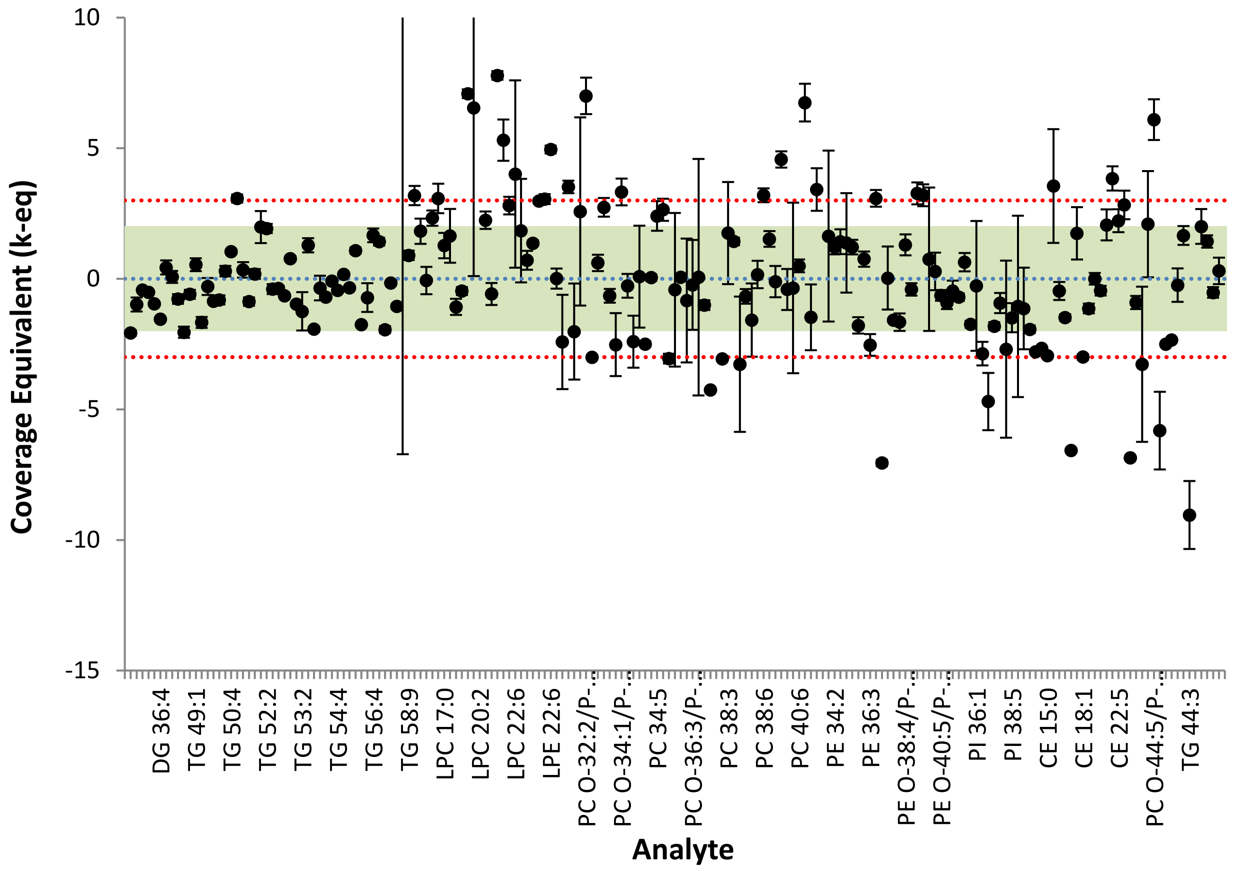

**Figure S6 SRM control for the presented workflow (RP-LC LILY): Accuracy assessment for SRM 1950 - "Metabolites in Frozen Human Plasma".**[8] Values are presented as normalized coverage equivalents at the mean (dots) and stdev (error bars, N=2) of measurements, overlaid onto the consensus mean value (blue line) and uncertainty (95% coverage-green region, 99% coverage-red region).

**Table S2 SRM control for the presented workflow (RP-LC LILY): Measurement accuracy summary compared against SRM 1950 - "Metabolites in Frozen Human Plasma"**

| Lipid Species | Measurement* | | | Consensus Value** | | | Units | No. of labs | Notes |
|---|---|---|---|---|---|---|---|---|---|
| DG 30:0 | 0.477 | ± | 0.018 | 0.83 | ± | 0.17 | nmol/mL | 7 | |
| DG 32:0 | 1.419 | ± | 0.322 | 2.6 | ± | 1.2 | nmol/mL | 11 | |
| DG 34:1 | 5.064 | ± | 0.118 | 6.1 | ± | 2.4 | nmol/mL | 16 | |
| DG 36:2 | 5.075 | ± | 0.059 | 6.2 | ± | 2.2 | nmol/mL | 16 | |
| DG 36:3 | 5.238 | ± | 0.092 | 8.4 | ± | 3.3 | nmol/mL | 15 | |
| DG 36:4 | 1.259 | ± | 0.017 | 2.8 | ± | 1.0 | nmol/mL | 12 | |
| TG 46:2 | 4.165 | ± | 0.349 | 3.6 | ± | 1.3 | nmol/mL | 8 | |
| TG 48:0 | 4.586 | ± | 0.275 | 4.5 | ± | 1.2 | nmol/mL | 10 | |
| TG 48:1 | 10.52 | ± | 0.588 | 13 | ± | 3.2 | nmol/mL | 16 | |
| TG 48:2 | 10.27 | ± | 0.599 | 16 | ± | 2.8 | nmol/mL | 15 | |
| TG 48:4 | 1.163 | ± | 0.04 | 1.3 | ± | 0.23 | nmol/mL | 5 | |
| TG 49:1 | 2.226 | ± | 0.103 | 2.0 | ± | 0.42 | nmol/mL | 9 | |
| TG 49:2 | 0.864 | ± | 0.117 | 1.8 | ± | 0.56 | nmol/mL | 6 | |
| TG 50:0 | 3.555 | ± | 0.269 | 3.8 | ± | 0.83 | nmol/mL | 11 | |
| TG 50:1 | 29.44 | ± | 0.937 | 38 | ± | 10 | nmol/mL | 14 | |
| TG 50:2 | 37.3 | ± | 2.091 | 47 | ± | 12 | nmol/mL | 15 | |
| TG 50:3 | 24.93 | ± | 1.322 | 23 | ± | 6.6 | nmol/mL | 16 | |
| TG 50:4 | 11.7 | ± | 0.251 | 8.7 | ± | 2.9 | nmol/mL | 15 | |
| TG 50:5 | 3.567 | ± | 0.1 | 1.6 | ± | 0.64 | nmol/mL | 7 | |
| TG 51:1 | 1.963 | ± | 0.142 | 1.8 | ± | 0.48 | nmol/mL | 7 | |
| TG 51:2 | 3.84 | ± | 0.176 | 4.8 | ± | 1.1 | nmol/mL | 8 | |
| TG 51:3 | 5.153 | ± | 0.358 | 4.8 | ± | 1.9 | nmol/mL | 5 | |
| TG 52:1 | 19.74 | ± | 1.78 | 14 | ± | 2.9 | nmol/mL | 11 | |
| TG 52:2 | 70.97 | ± | 2.458 | 44 | ± | 14 | nmol/mL | 16 | |
| TG 52:3 | 88.55 | ± | 5.513 | 100 | ± | 29 | nmol/mL | 16 | |
| TG 52:4 | 41.65 | ± | 1.823 | 48 | ± | 17 | nmol/mL | 15 | |
| TG 52:5 | 11.29 | ± | 0.188 | 15 | ± | 5.7 | nmol/mL | 13 | |
| TG 52:6 | 5.089 | ± | 0.129 | 4.0 | ± | 1.4 | nmol/mL | 8 | |
| TG 52:7 | 0.264 | ± | 0.009 | 0.39 | ± | 0.13 | nmol/mL | 5 | |
| TG 53:2 | 1.39 | ± | 0.301 | 1.9 | ± | 0.41 | nmol/mL | 9 | |
| TG 53:3 | 5.111 | ± | 0.299 | 3.7 | ± | 1.1 | nmol/mL | 6 | |
| TG 53:4 | 0.936 | ± | 0.072 | 2.4 | ± | 0.76 | nmol/mL | 6 | |
| TG 54:1 | 2.878 | ± | 0.43 | 3.2 | ± | 0.91 | nmol/mL | 10 | |
| TG 54:2 | 6.393 | ± | 0.141 | 8.2 | ± | 2.6 | nmol/mL | 13 | |
| TG 54:3 | 25.19 | ± | 0.708 | 26 | ± | 9.8 | nmol/mL | 15 | |
| TG 54:4 | 30.2 | ± | 0.96 | 36 | ± | 13 | nmol/mL | 15 | |
| TG 54:5 | 28.95 | ± | 0.64 | 27 | ± | 11 | nmol/mL | 15 | |
| TG 54:6 | 12.27 | ± | 0.219 | 14 | ± | 5.1 | nmol/mL | 16 | |

| | | | | | | | | |
|---|---|---|---|---|---|---|---|---|
| TG 54:7 | 7.223 | ± | 0.178 | 5.6 | ± | 1.5 | nmol/mL | 7 | |
| TG 56:2 | 0.286 | ± | 0.011 | 0.69 | ± | 0.23 | nmol/mL | 5 | |
| TG 56:3 | 1.299 | ± | 0.077 | 1.4 | ± | 0.14 | nmol/mL | 6 | |
| TG 56:4 | 2.931 | ± | 0.147 | 2.0 | ± | 0.56 | nmol/mL | 10 | |
| TG 56:5 | 6.089 | ± | 0.211 | 4.1 | ± | 1.4 | nmol/mL | 12 | |
| TG 56:7 | 7.746 | ± | 0.359 | 13 | ± | 2.7 | nmol/mL | 8 | |
| TG 56:9 | 0.668 | ± | 0.022 | 0.71 | ± | 0.27 | nmol/mL | 5 | |
| TG 58:7 | 1.321 | ± | 0.065 | 2.0 | ± | 0.64 | nmol/mL | 5 | |
| TG 58:8 | 14.66 | ± | 15.39 | 0.68 | ± | 0.21 | nmol/mL | 9 | |
| TG 58:9 | 1.443 | ± | 0.049 | 1.2 | ± | 0.27 | nmol/mL | 6 | |
| LPC 14:0 | 1.637 | ± | 0.074 | 1.0 | ± | 0.20 | nmol/mL | 16 | |
| LPC 15:0 | 0.72 | ± | 0.053 | 0.52 | ± | 0.11 | nmol/mL | 9 | |
| LPC 16:0 | 72.26 | ± | 5.76 | 73 | ± | 11 | nmol/mL | 20 | |
| LPC O-16:0 | 0.922 | ± | 0.047 | 0.55 | ± | 0.16 | nmol/mL | 10 | |
| LPC 16:1 | 3.477 | ± | 0.197 | 2.4 | ± | 0.35 | nmol/mL | 19 | |
| LPC 17:0 | 1.705 | ± | 0.117 | 1.4 | ± | 0.24 | nmol/mL | 6 | |
| LPC 18:0 | 32.43 | ± | 3.399 | 27 | ± | 3.3 | nmol/mL | 20 | |
| LPC 18:1 | 15.53 | ± | 0.71 | 18 | ± | 2.3 | nmol/mL | 19 | |
| LPC 18:2 | 20.64 | ± | 0.445 | 22 | ± | 2.9 | nmol/mL | 19 | |
| LPC 18:3 | 1.361 | ± | 0.022 | 0.44 | ± | 0.13 | nmol/mL | 18 | |
| LPC 20:1 | 0.347 | ± | 0.155 | 0.19 | ± | 0.024 | nmol/mL | 13 | |
| LPC 20:2 | 0.933 | ± | 0.016 | 0.23 | ± | 0.044 | nmol/mL | 9 | |
| LPC 20:3 | 2.383 | ± | 0.087 | 1.8 | ± | 0.26 | nmol/mL | 18 | |
| LPC 20:4 | 5.651 | ± | 0.253 | 6.0 | ± | 0.60 | nmol/mL | 20 | |
| LPC 20:5 | 1.046 | ± | 0.015 | 0.33 | ± | 0.092 | nmol/mL | 15 | |
| LPC 22:4 | 0.338 | ± | 0.032 | 0.12 | ± | 0.041 | nmol/mL | 8 | |
| LPC 22:5 | 0.794 | ± | 0.044 | 0.43 | ± | 0.13 | nmol/mL | 12 | |
| LPC 22:6 | 1.332 | ± | 0.502 | 0.77 | ± | 0.14 | nmol/mL | 17 | |
| LPE 16:0 | 1.408 | ± | 0.535 | 0.91 | ± | 0.27 | nmol/mL | 14 | |
| LPE 18:0 | 1.989 | ± | 0.199 | 1.6 | ± | 0.55 | nmol/mL | 15 | |
| LPE 18:1 | 2.042 | ± | 0.048 | 1.4 | ± | 0.47 | nmol/mL | 14 | |
| LPE 18:2 | 3.571 | ± | 0.049 | 1.9 | ± | 0.56 | nmol/mL | 16 | |
| LPE 20:4 | 2.351 | ± | 0.078 | 1.1 | ± | 0.41 | nmol/mL | 14 | |
| LPE 22:6 | 1.411 | ± | 0.027 | 0.52 | ± | 0.18 | nmol/mL | 12 | |
| PC 30:0 | 1.602 | ± | 0.123 | 1.6 | ± | 0.32 | nmol/mL | 11 | |
| PC 32:0 | 4.781 | ± | 1.805 | 7.2 | ± | 1.0 | nmol/mL | 18 | |
| PC O-32:0/31:0 | 2.942 | ± | 0.099 | 1.5 | ± | 0.41 | nmol/mL | 11 | Includes PC O-32:0 and PC [illegible] |
| PC 32:1 | 9.161 | ± | 3.495 | 13 | ± | 1.9 | nmol/mL | 18 | |
| PC O-32:1/P-32:0/31:1 | 2.219 | ± | 0.865 | 1.6 | ± | 0.24 | nmol/mL | 11 | Includes only PC P-32:0 a[illegible] |
| PC O-32:2/P-32:1/31:2 | 0.991 | ± | 0.065 | 0.34 | ± | 0.093 | nmol/mL | 8 | Includes only only PC P-3[illegible] |
| PC P-33:1/32:2 | 2.825 | ± | 0.116 | 2.6 | ± | 0.37 | nmol/mL | 16 | Includes only PC 32:2 res[illegible] |

| | | | | | | | | | |
|---|---|---|---|---|---|---|---|---|---|
| PC 34:0 | 3.112 | ± | 0.132 | 2.1 | ± | 0.37 | nmol/mL | 12 | |
| PC O-34:0/33:0 | 0.649 | ± | 0.045 | 0.76 | ± | 0.17 | nmol/mL | 10 | Includes PC O-34:0 and PC |
| PC 34:1 | 67.08 | ± | 25.32 | 120 | ± | 21 | nmol/mL | 19 | |
| PC O-34:1/P-34:0/33:1 | 7.759 | ± | 0.441 | 4.9 | ± | 0.86 | nmol/mL | 17 | Includes only PC P-34:0 an |
| PC O-34:2/P-34:1/33:2 | 4.851 | ± | 0.591 | 5.2 | ± | 1.3 | nmol/mL | 17 | Includes only PC P-34:1 an |
| PC O-34:3/P-34:2/33:3 | 2.581 | ± | 0.876 | 4.7 | ± | 0.88 | nmol/mL | 12 | Includes only PC P-34:2 an |
| PC P-35:1/34:2 | 243.8 | ± | 91.69 | 240 | ± | 47 | nmol/mL | 18 | Includes only PC 34:2 resu |
| PC P-35:2/34:3 | 7.753 | ± | 0.169 | 12 | ± | 1.7 | nmol/mL | 18 | Includes only PC 34:3 resu |
| PC O-35:4/34:4 | 1.011 | ± | 0.031 | 1.0 | ± | 0.25 | nmol/mL | 9 | Includes only PC 34:4 resu |
| PC 34:5 | 0.045 | ± | 0.003 | 0.034 | ± | 0.0045 | nmol/mL | 5 | |
| PC 36:1 | 38.15 | ± | 1.945 | 26 | ± | 4.6 | nmol/mL | 17 | |
| PC O-36:1/P-36:0/35:1 | 0.474 | ± | 0.186 | 3.5 | ± | 0.99 | nmol/mL | 16 | Includes only only PC 35:1 |
| PC 36:2 | 129.5 | ± | 73.51 | 140 | ± | 25 | nmol/mL | 18 | |
| PC O-36:2/P-36:1/35:2 | 7.495 | ± | 0.248 | 7.4 | ± | 1.7 | nmol/mL | 17 | Includes only PC P-36:1 an |
| PC 36:3 | 88.36 | ± | 33.2 | 100 | ± | 14 | nmol/mL | 17 | |
| PC O-36:3/P-36:2/35:3 | 3.508 | ± | 1.411 | 3.7 | ± | 0.82 | nmol/mL | 12 | Includes only PC P-36:2 an |
| PC O-36:4/P-36:3/35:4 | 12.09 | ± | 6.339 | 12 | ± | 1.4 | nmol/mL | 17 | Includes only PC P-36:3 an |
| PC 36:5 | 9.185 | ± | 0.334 | 11 | ± | 1.8 | nmol/mL | 16 | |
| PC O-36:5/P-36:4/35:5 | 0.099 | ± | 0.013 | 6.9 | ± | 1.6 | nmol/mL | 11 | Includes only only PC 35:5 |
| PC 38:2 | 8.372 | ± | 2.689 | 2.3 | ± | 0.20 | nmol/mL | 15 | |
| PC 38:3 | 35.1 | ± | 10.17 | 26 | ± | 5.2 | nmol/mL | 14 | |
| PC O-38:3/P-38:2/37:3 | 2.229 | ± | 0.074 | 1.5 | ± | 0.51 | nmol/mL | 14 | Includes only PC P-38:2 an |
| PC 38:4 | 38.2 | ± | 36.25 | 84 | ± | 14 | nmol/mL | 18 | |
| PC O-38:4/P-38:3/37:4 | 6.053 | ± | 0.56 | 7.4 | ± | 2.0 | nmol/mL | 12 | Includes only PC P-38:3 an |
| PC 38:5 | 29.52 | ± | 11.07 | 42 | ± | 7.9 | nmol/mL | 18 | |
| PC O-38:5/P-38:4/37:5 | 11.26 | ± | 0.84 | 11 | ± | 1.6 | nmol/mL | 16 | Includes only only PC P-38 |
| PC 38:6 | 55.07 | ± | 1.17 | 41 | ± | 4.4 | nmol/mL | 18 | |
| PC O-38:6/P-38:5/37:6 | 5.125 | ± | 0.301 | 3.6 | ± | 1.0 | nmol/mL | 12 | Includes only PC P-38:5 an |
| PC P-38:6/36:0 | 1.158 | ± | 0.234 | 1.2 | ± | 0.39 | nmol/mL | 10 | Includes PC P-38:6 and PC |
| PC 40:4 | 4.59 | ± | 0.115 | 2.9 | ± | 0.37 | nmol/mL | 18 | |
| PC O-40:4/P-40:3/39:4 | 0.794 | ± | 0.298 | 0.95 | ± | 0.38 | nmol/mL | 8 | Includes only only PC P-40 |
| PC 40:5 | 6.314 | ± | 3.592 | 6.7 | ± | 1.1 | nmol/mL | 18 | |
| PC 40:6 | 15.29 | ± | 0.638 | 14 | ± | 2.6 | nmol/mL | 17 | |
| PC 40:7 | 5.254 | ± | 0.188 | 3.5 | ± | 0.26 | nmol/mL | 16 | |
| PC O-40:7/P-40:6/39:7 | 0.761 | ± | 0.29 | 1.1 | ± | 0.23 | nmol/mL | 9 | Includes only only PC P-40 |
| PC 40:8 | 1.414 | ± | 0.163 | 0.73 | ± | 0.20 | nmol/mL | 14 | |
| PC O-42:5/P-42:4 | 3.009 | ± | 0.126 | 0.79 | ± | 0.12 | nmol/mL | 7 | Includes only PC P-42:4 re |
| PE 34:1 | 1.478 | ± | 0.556 | 1.2 | ± | 0.17 | nmol/mL | 14 | |
| PE 34:2 | 2.514 | ± | 0.071 | 2.2 | ± | 0.26 | nmol/mL | 16 | |
| PE O-34:2/P-34:1 | 1.021 | ± | 0.081 | 0.78 | ± | 0.17 | nmol/mL | 11 | Includes only PE P-34:1 re |
| PE O-34:3/P-34:2 | 2.064 | ± | 0.781 | 1.5 | ± | 0.41 | nmol/mL | 11 | Includes only PE P-34:2 re |

| | | | | | | | | |
|---|---|---|---|---|---|---|---|---|
| PE 36:1 | 1.615 | ± | 0.073 | 1.3 | ± | 0.26 | nmol/mL | 14 | |
| PE 36:2 | 5.29 | ± | 0.247 | 6.7 | ± | 0.79 | nmol/mL | 16 | |
| PE O-36:2/P-36:1/35:2 | 1.096 | ± | 0.064 | 0.93 | ± | 0.22 | nmol/mL | 12 | Includes only only PE P-36 |
| PE 36:3 | 1.437 | ± | 0.158 | 2.4 | ± | 0.38 | nmol/mL | 16 | |
| PE O-36:3/P-36:2/35:3 | 5.541 | ± | 0.243 | 3.2 | ± | 0.76 | nmol/mL | 15 | Includes only only PE P-36 |
| PE 36:4 | 0.35 | ± | 0.049 | 3.1 | ± | 0.39 | nmol/mL | 16 | |
| PE O-36:4/P-36:3 | 1.609 | ± | 0.35 | 1.6 | ± | 0.29 | nmol/mL | 14 | Includes only PE P-36:3 re |
| PE O-36:5/P-36:4 | 1.9 | ± | 0.076 | 4.9 | ± | 1.9 | nmol/mL | 15 | Includes only PE P-36:4 re |
| PE 38:4 | 6.106 | ± | 0.401 | 8.1 | ± | 1.2 | nmol/mL | 16 | |
| PE O-38:4/P-38:3/37:4 | 1.174 | ± | 0.072 | 0.94 | ± | 0.18 | nmol/mL | 9 | Includes only only PE P-38 |
| PE 38:5 | 2.509 | ± | 0.112 | 2.7 | ± | 0.47 | nmol/mL | 12 | |
| PE O-38:5/P-38:4 | 12.01 | ± | 0.798 | 5.8 | ± | 1.9 | nmol/mL | 17 | Includes only PE P-38:4 re |
| PE 38:6 | 5.087 | ± | 0.248 | 3.2 | ± | 0.59 | nmol/mL | 15 | |
| PE O-38:6/P-38:5 | 5.799 | ± | 3.295 | 4.9 | ± | 1.2 | nmol/mL | 16 | Includes only PE P-38:5 re |
| PE O-38:7/P-38:6 | 3.777 | ± | 0.705 | 3.5 | ± | 0.98 | nmol/mL | 8 | Includes only PE P-38:6 re |
| PE O-40:5/P-40:4/39:5 | 0.647 | ± | 0.027 | 0.73 | ± | 0.13 | nmol/mL | 12 | Includes only only PE P-40 |
| PE 40:6 | 1.467 | ± | 0.084 | 1.8 | ± | 0.36 | nmol/mL | 14 | |
| PE O-40:6/P-40:5/39:6 | 1.156 | ± | 0.122 | 1.3 | ± | 0.31 | nmol/mL | 14 | Includes only only PE P-40 |
| PE O-40:7/P-40:6/39:7 | 1.994 | ± | 0.104 | 2.5 | ± | 0.72 | nmol/mL | 14 | Includes only only PE P-40 |
| PI 34:1 | 2.667 | ± | 0.148 | 2.4 | ± | 0.42 | nmol/mL | 14 | |
| PI 34:2 | 2.138 | ± | 0.05 | 2.8 | ± | 0.38 | nmol/mL | 14 | |
| PI 36:1 | 1.939 | ± | 1.467 | 2.1 | ± | 0.59 | nmol/mL | 13 | |
| PI 36:2 | 5.035 | ± | 0.42 | 7.7 | ± | 0.93 | nmol/mL | 15 | |
| PI 36:3 | 0.837 | ± | 0.318 | 2.2 | ± | 0.29 | nmol/mL | 14 | |
| PI 36:4 | 2.132 | ± | 0.07 | 3.0 | ± | 0.48 | nmol/mL | 14 | |
| PI 38:3 | 2.898 | ± | 0.208 | 3.4 | ± | 0.54 | nmol/mL | 14 | |
| PI 38:4 | 13.07 | ± | 7.459 | 19 | ± | 2.2 | nmol/mL | 17 | |
| PI 38:5 | 1.844 | ± | 0.245 | 2.5 | ± | 0.44 | nmol/mL | 15 | |
| PI 40:4 | 0.256 | ± | 0.146 | 0.30 | ± | 0.042 | nmol/mL | 7 | |
| PI 40:6 | 0.658 | ± | 0.25 | 0.84 | ± | 0.16 | nmol/mL | 12 | |
| PG 34:1 | 0.141 | ± | 0.086 | 1.3 | ± | 0.60 | nmol/mL | 5 | |
| CE 16:0 | 416 | ± | 126.3 | 210 | ± | 58 | nmol/mL | 13 | |
| CE 16:1 | 87.4 | ± | 9.303 | 100 | ± | 27 | nmol/mL | 11 | |
| CE 16:2 | 1.219 | ± | 0.057 | 1.9 | ± | 0.46 | nmol/mL | 5 | |
| CE 17:1 | 1.629 | ± | 0.078 | 8.2 | ± | 1.0 | nmol/mL | 9 | |
| CE 18:0 | 21.44 | ± | 3.708 | 15 | ± | 3.7 | nmol/mL | 7 | |
| CE 18:1 | 120.8 | ± | 11.99 | 450 | ± | 110 | nmol/mL | 14 | |
| CE 18:2 | 1211 | ± | 80.09 | 1,700 | ± | 430 | nmol/mL | 14 | |
| CE 18:3 | 84 | ± | 5.329 | 84 | ± | 24 | nmol/mL | 13 | |
| CE 20:3 | 29.54 | ± | 2.224 | 35 | ± | 12 | nmol/mL | 13 | |
| CE 20:4 | 469.7 | ± | 34.45 | 350 | ± | 58 | nmol/mL | 14 | |

| | | | | | | | | | |
|---|---|---|---|---|---|---|---|---|---|
| CE 20:5 | 71.04 | ± | 3.981 | 38 | ± | 8.6 | nmol/mL | 12 | |
| CE 22:5 | 7.657 | ± | 0.702 | 4.1 | ± | 1.6 | nmol/mL | 6 | |
| CE 22:6 | 63.86 | ± | 5.203 | 37 | ± | 9.5 | nmol/mL | 11 | |
| Cholesterol | 16.43 | ± | 1.209 | 770 | ± | 110 | nmol/mL | 8 | |
| LPC O-18:1 | 0.292 | ± | 0.033 | 0.41 | ± | 0.13 | nmol/mL | 3 | |
| PC 28:0 | 0.078 | ± | 0.074 | 0.16 | ± | 0.025 | nmol/mL | 4 | |
| PC O-42:6/P-42:5 | 0.753 | ± | 0.285 | 0.46 | ± | 0.14 | nmol/mL | 4 | Includes only PC P-42:5 re |
| PC O-44:5/P-44:4 | 3.128 | ± | 0.234 | 1.3 | ± | 0.30 | nmol/mL | 3 | Includes only PC P-44:4 re |
| PE 34:3 | 0.024 | ± | 0.03 | 0.14 | ± | 0.020 | nmol/mL | 4 | |
| PS 36:2 | 0.342 | ± | 1.025 | 0.73 | ± | 1.6 | nmol/mL | 4 | |
| TG 42:0 | 0.695 | ± | 0.069 | 0.38 | ± | 0.19 | nmol/mL | 5 | |
| TG 44:3 | 0.095 | ± | 0.012 | 0.18 | ± | 0.0094 | nmol/mL | 4 | |
| TG 47:2 | 0.659 | ± | 0.044 | 0.21 | ± | 0.027 | nmol/mL | 3 | |
| TG 49:0 | 0.42 | ± | 0.037 | 0.31 | ± | 0.055 | nmol/mL | 3 | |
| TG 54:8 | 1.349 | ± | 0.071 | 0.92 | ± | 0.30 | nmol/mL | 4 | |
| TG 55:3 | 0.352 | ± | 0.03 | 0.43 | ± | 0.15 | nmol/mL | 3 | |
| TG 58:11 | 0.115 | ± | 0.009 | 0.11 | ± | 0.018 | nmol/mL | 3 | |

* Measurement mean ± 1

** Consensus mean ± sta

### 8. Bibliography

(1) Neubauer, S.; Haberhauer-Troyer, C.; Klavins, K.; Russmayer, H.; Steiger, M. G.; Gasser, B.; Sauer, M.; Mattanovich, D.; Hann, S.; Koellensperger, G. U13C Cell Extract of Pichia Pastoris - A Powerful Tool for Evaluation of Sample Preparation in Metabolomics. *J. Sep. Sci.* **2012**, *35* (22), 3091–3105. https://doi.org/10.1002/jssc.201200447.

(2) Schoeny, H.; Rampler, E.; Hermann, G.; Grienke, U.; Rollinger, J. M.; Koellensperger, G. Preparative Supercritical Fluid Chromatography for Lipid Class Fractionation — a Novel Strategy in High-Resolution Mass Spectrometry Based Lipidomics. *Anal. Bioanal. Chem.* **2020**, *412*, 2365–2374.

(3) Matyash, V.; Liebisch, G.; Kurzchalia, T. V.; Shevchenko, A.; Schwudke, D. Lipid Extraction by Methyl- *Tert* -Butyl Ether for High-Throughput Lipidomics. *J. Lipid Res.* **2008**, *49* (5), 1137–1146. https://doi.org/10.1194/jlr.D700041-JLR200.

(4) Lipidomics Standards Initiativ. Lipid Species Quantifciaton https://lipidomics-standards-initiative.org/guidelines/lipid-species-quantification (accessed Feb 12, 2020).

(5) Wang, M.; Wang, C.; Han, X. Selection of Internal Standards for Accurate Quantification of Complex Lipid Species in Biological Extracts by Electrospray Ionization Mass Spectrometry—What, How and Why? *Mass Spectrom. Rev.* **2017**, *36* (6), 693–714. https://doi.org/DOI 10.1002/mas.21492.

(6) Triebl, A.; Burla, B.; Selvalatchmanan, J.; Oh, J.; Tan, S. H.; Chan, M. Y.; Mellet, N. A.; Meikle, P. J.; Torta, F.; Wenk, M. R. Shared Reference Materials Harmonize Lipidomics across MS-Based Detection Platforms and Laboratories. *J. Lip* **2020**, *61*, 105–115. https://doi.org/10.1194/jlr.D119000393.

(7) Food and Drug Administration. *Bioanalytical Method Validation Guidance for Industry*; 2018.

(8) Ulmer, C. Z.; Ragland, J. M.; Koelmel, J. P.; Heckert, A.; Jones, C. M.; Garrett, T.; Yost, R. A.; Bowden, J. A. LipidQC: Method Validation Tool for Visual Comparison to SRM 1950 Using NIST Interlaboratory Comparison Exercise Lipid Consensus Mean Estimate Values. *Anal. Chem.* **2017**, *89*, 13069–13073. https://doi.org/10.1021/acs.analchem.7b04042.
